# Glucosylceramide induced ectosomes propagate pathogenic α-synuclein in Parkinson’s disease

**DOI:** 10.1101/2025.02.14.638390

**Authors:** Julie Jacquemyn, Brian Mariott, Jinlan Chang, Nathanael Y J Lee, Luis F. Rubio Atonal, Ceili Green, Jeremy Wong, Kennedi Chik, Claudia Acevedo-Morantes, Carol X.-Q. Chen, Michael Nicouleau, Zhipeng You, Eric Deneault, Narges Abdian, Thomas M. Durcan, Jesse Jackson, Maria S. Ioannou

## Abstract

Intercellular transmission of α-synuclein contributes to Parkinson’s disease pathology. Yet, the mechanisms of α-synuclein spread are not fully understood. Here, we used live-cell microscopy to examine the impact of Parkinson’s disease associated lipid alterations on α-synuclein release. We discovered that increased glucosylceramides induce ectosome shedding from primary neurons, and from dopaminergic neurons derived from Parkinson’s disease patient iPSCs harboring mutations in *GBA1* (N370S, L444P and W378G) and *LRRK2* (G2019S and R1441H) compared to their isogenic control. We show that elevated glucosylceramide similarly increases vesicle release and uptake by other neurons in living mouse brains using 2-photon microscopy. Finally, we show that ectosomes are loaded with pathogenic α-synuclein and lead to the transmission of α-synuclein pathology to neighbouring neurons. These data reveal ectosomes as the predominant route for α-synuclein transmission that can only be appreciated by live-cell imaging technologies.

## INTRODUCTION

Intercellular transmission of misfolded α-synuclein has been proposed to play a key role in Parkinson’s disease (PD)^1,2^. Dysregulated lipid metabolism is also a common feature of PD, although its contribution to disease pathology is less clear^3–5^. The most common genetic risk factors for PD are mutations in *GBA1*^6–8^, a gene which encodes the sphingolipid metabolism enzyme β-glucocerebrosidase (GCase). GCase hydrolyzes glucosylceramide to ceramide and glucose^9^. *L*oss-of-function *GBA1* mutations lead to reduced GCase activity, increased glucosylceramide levels and subsequent accumulation of pathological α-synuclein contributing to a toxic cascade^10–16^. These features, reduced GCase activity and increased glucosylceramide levels, are not specific to *GBA1-*PD patients as they also occur in familial PD caused by mutations in the leucine-rich repeat kinase 2 (*LRRK2*) and sporadic PD^10,13,17–21^. This suggests that dysregulated sphingolipid metabolism could drive PD pathology. But how this contributes to the prion-like transmission of α-synuclein pathology between neurons is poorly understood.

One possibility for how glucosylceramide alterations could affect the transmission of α-synuclein is through extracellular vesicles^22^. While extracellular vesicles have been proposed as a vehicle for the transmission of pathogenic proteins, the use of biochemical purification precludes the ability to discern the identity of extracellular vesicles^23^. Due to overlapping size and a lack of reliable markers, extracellular vesicle subtypes cannot be distinguished from one another once released^23^. With different mechanisms of formation and function, these vesicle subtypes need to be studied independently to gain a comprehensive understanding of their role in PD.

Ectosomes are a subtype of extracellular vesicles that form by the outward budding and scission of the plasma membrane^23^. Lipid rafts, which serve as a platform for ectosome biogenesis^24^, are enriched in PD-elevated sphingolipids. α-Synuclein is present in the cytosol and interacts with these raft-associated lipids on the plasma membrane^25–28^. These PD-associated sphingolipids promote the formation of oligomeric α-synuclein species associated with PD pathology^29^. This raises the possibility that ectosomes contribute to the intercellular transmission of α-synuclein. However, because ectosomes can only be accurately identified in the act of release by imaging techniques, they have been largely understudied. Here, using multiple live-cell and super-resolution imaging modalities, we discovered that reduced GCase activity and increased glucosylceramide levels stimulate ectosome shedding from primary cortical neurons, iPSC-derived dopaminergic neurons from patients with mutations in *GBA1* (N370S, L444P or W378G) and *LRRK2* (G2019S or R1441H), and from cortical neurons in vivo. These ectosomes serve as a major and previously unappreciated route for the intercellular transport of misfolded α-synuclein thereby propagating pathology in PD.

## RESULTS

### Glucosylceramide stimulates ectosome shedding from primary neurons

Glucosylceramide is synthesized and secreted by neurons in extracellular vesicles^30^. Since glucosylceramide is enriched on the plasma membrane^24^ and is altered in PD^10,17–20,31^, we explored if glucosylceramide affects ectosome shedding. In PD, cortical neurons develop α-synuclein inclusions and are affected at late stages of the disease^32,33^. The plasma membrane of primary rat cortical neurons was labelled by expressing mCherry attached to the C-terminal domain of CD58 that binds glycosylphosphatidylinositol (GPI), anchoring it to the membrane. This construct is herein referred to as mCh-GPI. Neurons were treated with glucosylceramide tagged with the fluorophore nitrobenzofurazan (NBD) and imaged live. As expected, NBD-glucosylceramide is highly enriched on the plasma membrane and induced a ∼26-fold increase in vesicle budding from the plasma membrane compared to control neurons (Fig. 1a-b). The plasma membrane vesicles bud from thick neurites and the soma (Fig. 1a and Extended Data Fig. 1a). This is distinct from neuritic beading which occurs along constricted neurites of dying cells and can also give rise to extracellular vesicles^24^. Consistently, no changes in non-specific cell death were detected with NBD-glucosylceramide treatment as measured with ethidium homodimer-1 staining (Extended Data Fig. 1b). Nor did NBD-glucosylceramide treatment affect cytotoxicity as assessed using an MTT assay (Extended Data Fig. 1c). This indicates these plasma membrane vesicles are ectosomal in identity (also known as microvesicles). Performing time-lapse imaging, we see the plasma membrane bud grows outward as the neck region constricts, the vesicle detaches and often disappears from the field of view (Fig. 1c and Extended Data Fig. 1d). These vesicle buds are the same diameter as detached vesicles in the media (1 to 2.5 µm) suggesting they are being released (Fig. 1d-f). Several ectosomal buds and detached vesicles contain smaller vesicles within (Extended Data Fig. 1e), however the physiological relevance of these intraluminal ectosomes remains unclear^34,35^.

**Fig. 1.**
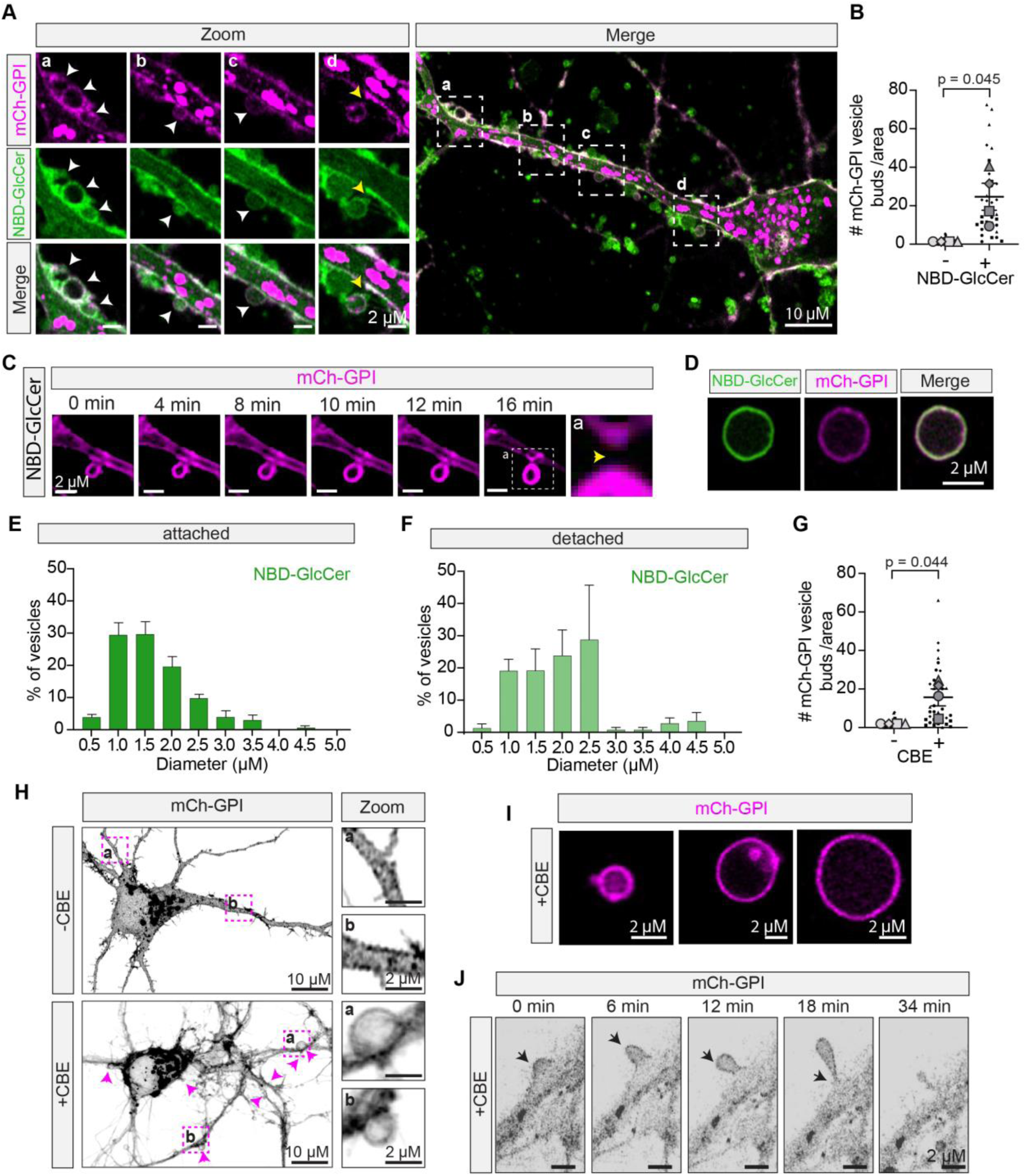
Exogenous NBD-GlcCer or GCase inhibition induce ectosome formation. **(A)** Live-cell image of cortical neuron expressing mCh-GPI treated with NBD-glucosylceramide (GlcCer). Boxes highlight ectosomes forming from the plasma membrane and are magnified on the left. White arrows indicate ectosomes forming. Yellow arrow indicates ectosome detached from neurite. **(B)** Amount of mCh-GPI ectosome buds per cell area relative to control. *n* = 4 independent experiments; 10 cells/coverslip/treatment; mean ± SEM; One sample t-test. Graph is depicted as a superplot where biological replicates are shown in large shapes and technical replicates are shown as small shapes. **(C)** Time-lapse imaging showing mCh-GPI-positive ectosome with NBD-GlcCer treatment. **(D)** Image of mCh-GPI-positive vesicle from neuron conditioned media after NBD-GlcCer treatment. (**E-F**) Size distribution of attached and detached mCh-GPI-positive vesicles from NBD-GlcCer treated neurons. *n* = 4 independent experiments; bars show mean ± SEM. **(G)** Amount of mCh-GPI ectosome buds per cell area relative to control. *n* = 4 independent experiments; 10 cells/coverslip/treatment; mean ± SEM; One sample t-test. Graph is depicted as a superplot where biological replicates are shown in large shapes, and technical replicates are shown as small shapes. **(H)** Live-cell image of cortical neuron expressing mCh-GPI treated with 100 µM CBE. The arrows highlight vesicles budding from the plasma membrane along the neurites and soma. Boxes show magnified images on the right. **(I)** Image of mCh-GPI-positive vesicles from neuron-conditioned media with CBE treatment. **(J)** Time-lapse series of vesicle formation, extension and shedding in primary cortical neurons with the addition of 100 µM CBE. Black arrows indicate vesicle formation over time.

Since glucosylceramide levels increase with decreased GCase activity in familial and sporadic PD^10–12,14,15,17–21,31^, we next tested whether inhibiting GCase with the selective small molecule inhibitor conduritol-β-epoxide (CBE)^36^ affects ectosome formation. We confirmed the ability of CBE to reduce GCase activity (Extended Data Fig. 1f) and accumulate glucosylceramide (Fig. Extended Data Fig. 1g-h) in cortical neurons. Similar to adding glucosylceramide directly, CBE also stimulated mCh-GPI positive ectosome budding from the soma and neurites of cortical neurons (Fig. 1g-h). These plasma membrane buds have a similar diameter (1 to 2.5 µm) to vesicles in the media (Fig. 1i and Extended Data Fig. 1i-j), further supporting their release. Time-lapse imaging reveals that these ectosomes form by the outward budding and fission of the plasma membrane in response to CBE treatment (Fig. 1j). No changes in non-specific cell death or cytotoxicity were observed with CBE treatment by ethidium homodimer-1 (Extended Data Fig. 1k), or MTT assay (Extended Data Fig. 1l), respectively. Collectively, our data indicate that decreased GCase activity and increased glucosylceramide induce the formation of ectosomes from cortical rat neurons.

### Glucosylceramide stimulates vesicle shedding from neurons in vivo

Elevated glucosylceramide levels induce ectosome formation in vitro, prompting us to investigate whether this also happens in vivo. To image vesicle formation in a living brain using two-photon microscopy requires neurons to be labelled with a fluorophore that is present in the soma, neurites and ectosomes. Since ectosomes contain cytosolic proteins^37,38^, we reasoned that expression of the soluble fluorophore tdTomato could be used to follow their formation. We first tested this concept by expressing tdTomato in our primary rat cortical neurons. Live-cell imaging confirmed that ∼60% of the ectosomes contained tdTomato after addition of NBD-glucosylceramide (Fig. 2a and Extended Data Fig. 2a). TdTomato is also present within vesicles in the media (Fig. 2b), indicating these vesicles are released. This data validates the use of tdTomato to visualize ectosome budding by live-cell imaging.

**Fig. 2.**
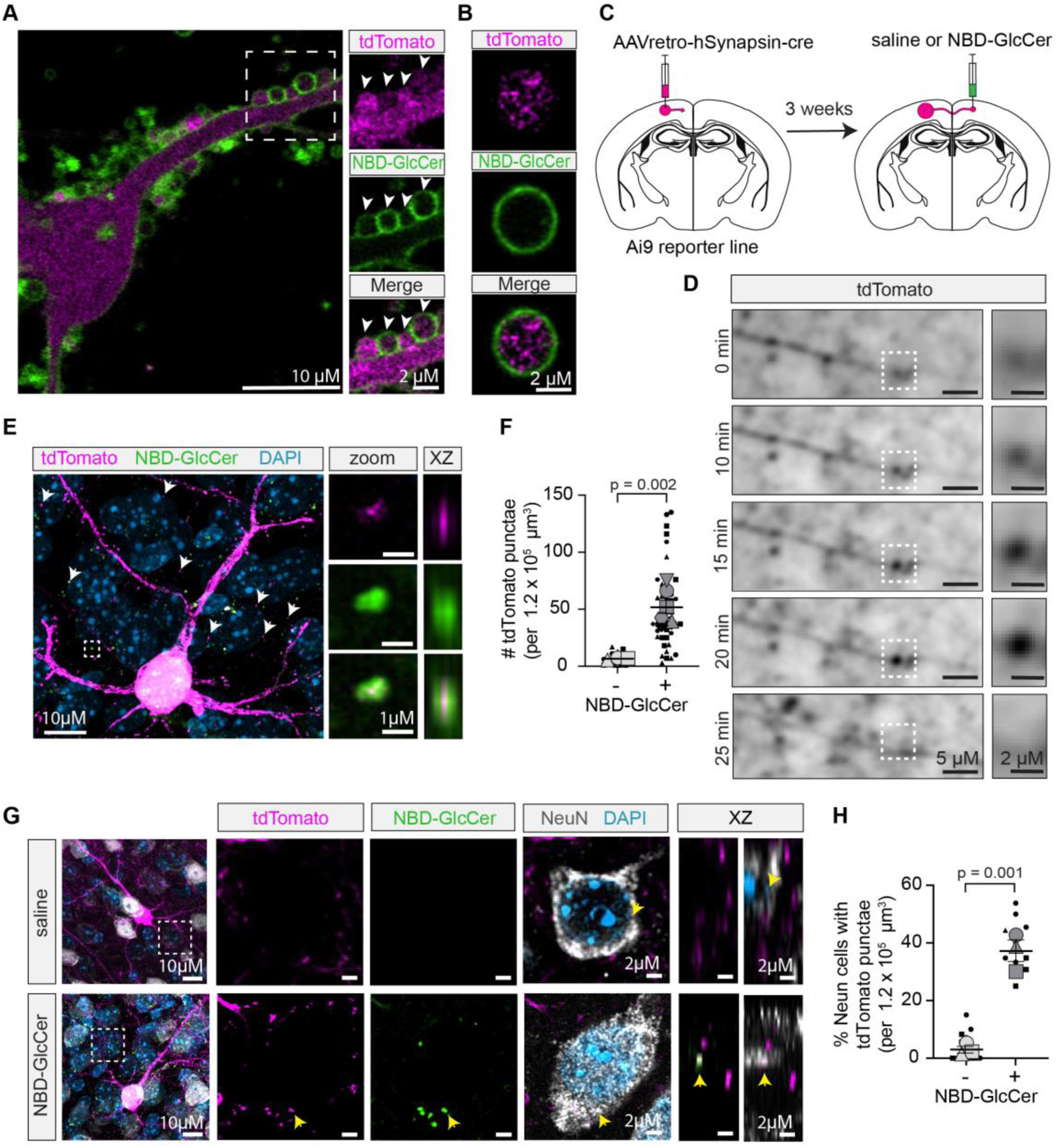
Glucosylceramide-induced ectosome formation in an in vivo mouse brain. **(A)** Live-cell image of cortical neuron expressing tdTomato treated with NBD-GlcCer. The boxed areas highlighting vesicles containing tdTomato and are magnified on the right. **(B)** Image of NBD-GlcCer-positive vesicle containing tdTomato from neuron conditioned media. **(C)** Approach to sparsely label cortical neurons with tdTomato in vivo where NBD-GlcCer was later injected. **(D)** Two-photon time-lapse imaging of the cortex of an anesthetized mouse showing a mCh-GPI-positive vesicle growing along the neurite and disappearing. Boxed areas are magnified on the right. **(E)** Maximum intensity projection of fixed tissue showing a cortical neuron expressing tdTomato after NBD-GlcCer administration. Arrows indicate several tdTomato-positive vesicles detached from neuron. Boxed area shows magnification of example vesicle with orthogonal views on the right confirming it is detached. **(F)** tdTomato-positive puncta ≥ 2 µM from tdTomato-positive neuron. *n* ≥ 3 animals; 5 images per n; mean ± SEM; Unpaired t-test. Graph is depicted as a superplot where biological replicates are shown in large shapes, and technical replicates are shown as small shapes. **(G)** Maximum intensity projection showing a cortical neuron expressing tdTomato after NBD-GlcCer administration and immunostained for NeuN. Boxed area, single slice magnification and orthogonal views showing tdTomato-positive puncta internalized by a neighboring neuron stained with NeuN in the presence of NBD-GlcCer. **(H)** Percentage of NeuN positive neurons neighboring the tdTomato-expressing neuron containing tdTomato puncta. *n* = 3 animals; 5 images per n; mean ± SEM; Unpaired t-test. Graph is depicted as a superplot where biological replicates are shown in large shapes, and technical replicates are shown as small shapes.

For in vivo experiments, the cortex was targeted because of its accessibility by two-photon microscopy and because PD pathology including α-synuclein inclusions spread to the cortex at the late stages of the disease^32,33^. To visualize ectosomes forming from neurons in vivo, sparse labelling of the tdTomato marker is required. To achieve this, we injected a retrograde AAV expressing cre under the synapsin promoter in the motor cortex of tdTomato reporter mice. This led to sparse labelling of cortical neurons in the contralateral hemisphere, where we then injected NBD-glucosylceramide or saline and imaged using two-photon microscopy in anesthetized mice (Fig. 2c and Extended Data Fig. 2b). Similar to our in vitro experiments, we observed tdTomato-positive vesicles growing from a neurite that quickly disappeared (Fig. 2d). To investigate this further, we analyzed fixed tissue using super-resolution microscopy to quantify detached tdTomato-positive puncta within a 3D volume. To avoid the possibility of counting dendritic spines, we quantified the number of tdTomato-positive puncta at least 2 µm away from a tdTomato-positive neuron. We discovered a significant increase in detached tdTomato puncta upon the addition of NBD-glucosylceramide compared to saline controls (Fig. 2e-f and Extended Data Fig. 2c). Several tdTomato-puncta were also positive for NBD-glucosylceramide (Fig. 2e).

Since propagation of pathogenic proteins occurs via intercellular transport between neurons, we next wondered whether these tdTomato positive puncta could be taken up by neighboring neurons. Although it is not possible to ascertain the identity of vesicle subtype once taken up by another cell, surrounding neurons contained more tdTomato positive puncta upon addition of NBD-glucosylceramide compared to the saline control (Fig. 2g-h, Extended Data Fig. 2d-e, and Supplementary Video 1). TdTomato puncta inside the neurons were also positive for NBD-glucosylceramide (Extended Data Fig. 2f). Overall, this data is consistent with glucosylceramide induced vesicle formation from neurons in vivo, with the potential to spread to neighboring neurons. To directly explore the role of ectosomes in the process, we turned back to our in vitro model system.

### iPSC-derived neurons with *GBA1* and *LRRK2* mutations shed more ectosomes

Glucosylceramide and reduced GCase activity induce ectosome shedding and both are observed in PD caused by mutations in *GBA1* and *LRRK2*^10–12,14,15,17–21,31^. Thus, we next sought to determine if PD neurons also shed more ectosomes. We generated neurons with approximately 70% dopaminergic identity (Extended Data Fig. 3a-b) from PD patient iPSC lines with mutations in *GBA1* (N370S, L444P or W378G), *LRRK2* (G2019S or R1441H), and their corresponding isogenic controls^39,40^. Loss of midbrain dopaminergic neurons is an early and prominent feature of PD making dopaminergic neuron cultures a highly relevant model system to study the disease^41,42^. Consistent with previous work^10–12,14^, dopaminergic neurons with mutations in *GBA1* and *LRRK2* have reduced GCase activity compared to their isogenic controls (Fig. 3a).

**Fig. 3.**
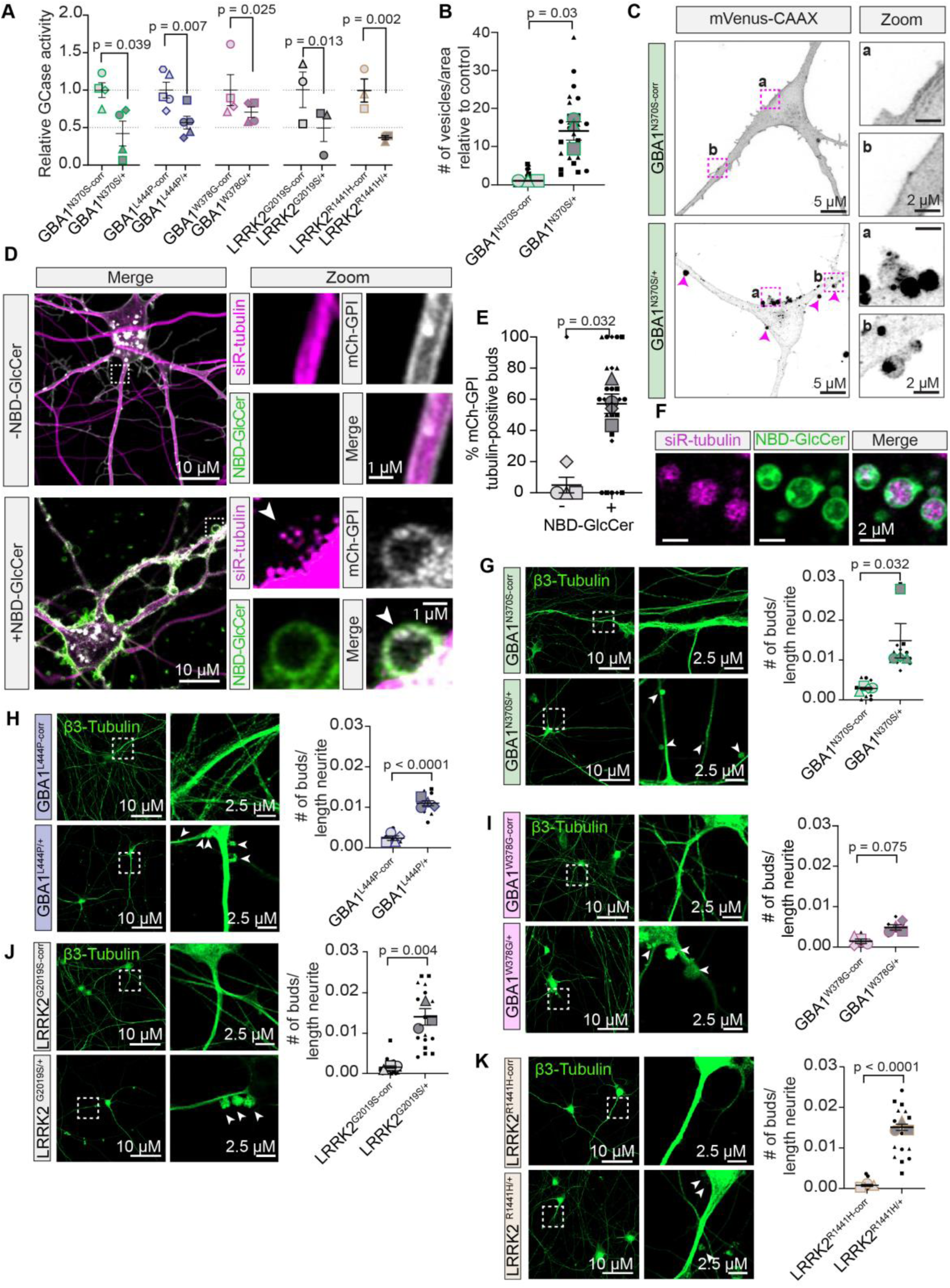
Dopaminergic neurons derived from PD-patient iPSCs shed more ectosomes. **(A)** GCase activity of dopaminergic neurons normalized to their isogenic corrected control (corr) from 3 or more independent inductions per line; mean ± SEM; One sample t-test. Graph shows experimental averages. **(B)** Number of mVenus-CAAX positive ectosome buds per cell area in *GBA1*^N370S/+^ and control dopaminergic neuron. *n* = 3 independent experiments; 10 cells per experiment; mean ± SEM; Mann-Whitney test. Graph is depicted as a superplot where biological replicates are shown in large shapes, and technical replicates are shown as small shapes. **(C)** Live-cell maximum intensity projection of *GBA1*^N370S/+^ and control neurons expressing mVenus-CAAX. Boxed areas highlighting ectosomes forming from plasma membrane magnified on the right. **(D)** Live-cell image of cortical neurons expressing mCh-GPI treated with siR-tubulin and NBD-GlcCer. Boxed areas showing tubulin within ectosome bud are magnified on the right. **(E)** Percentage of mVenus-CAAX positive buds containing siR-tubulin induced by NBD-GlcCer. *n* = 3 independent experiments; 10 cells per experiment; mean ± SEM; Mann-Whitney test. Graph is depicted as a superplot where biological replicates are shown in large shapes, and technical replicates are shown as small shapes. **(F)** Image of vesicles containing siR-tubulin from neuronal conditioned media after NBD-GlcCer treatment. (**G-K**) Tile scan of fixed iPSC-derived dopaminergic neurons from *GBA1* and *LRRK2* mutant lines and their isogenic controls stained for β3-tubulin. Magnified images on the right. Arrows indicate tubulin-positive vesicles budding from the neurite. Quantification of tubulin-positive buds relative to control. *n* ≥ 3 independent experiments; 3 tile scans/ 2 coverslip/experiment. Mean ± SEM; Unpaired t-test. Graph is depicted as a superplot where biological replicates are shown in large shapes, and technical replicates are shown as small shapes.

By live-cell imaging of the plasma membrane marker mVenus-CAAX, we found a ∼15-fold increase in ectosomes shedding from *GBA1*-N370S and *LRRK2*-G2019S neurons compared to their isogenic controls (Fig. 3b-c and Extended Data Fig. 3c-d). Additionally, these plasma membrane buds have a similar diameter (1 to 2.5 µm) to CBE and NBD-GlcCer-induced vesicles (Extended Data Fig. 3e-f). Toxicity, as determined by MTT assay, was not affected by the expression of mVenus-CAAX, or in *GBA1*-N370S relative to their isogenic control (Extended Data Fig. 3g-h). There was a slight but non-significant increase in toxicity in *LRRK2*-G2019S mutants relative to their isogenic control (Extended Data Fig. 3h). To make sure overexpression of the membrane targeting proteins are not required for inducing ectosomes, we assessed additional markers associated with ectosomes. The microtubule-associated protein tau is enriched in large vesicles that are proposed to be ectosomes^38^. Using differential ultracentrifugation to separate vesicles based on size (Extended Data Fig. 4a), we confirmed increased release of large vesicles containing the extracellular vesicle marker CD81 (Extended Data Fig. 4b-c) and tubulin (Extended Data Fig. 4b,d) in *GBA1*-N370S neurons compared to their isogenic controls. We next confirmed the presence of siR-tubulin in ectosomes budding from CBE- and NBD-glucosylceramide treated cortical neurons and *GBA1*-N370S dopaminergic neurons (Fig. 3d-e and Extended Data Fig. 4e-f). 60% and 70% of the ectosome buds treated with NBD-glucosylceramide and CBE respectively, contained tubulin (Fig. 3e and Extended Data Fig. 4g). Even more ectosomes, 80%, from the *GBA1*-N370S dopaminergic neurons contained tubulin (Fig. Extended Data Fig. 4h). Tubulin-positive vesicle buds have a similar diameter to those described above (Fig. 1d, Extended Data Fig. 4i-j). SiR-tubulin is also present within vesicles released into the neuron-conditioned media (Fig. 3f). siR-tubulin did not affect cytotoxicity in CBE- and NBD-glucosylceramide treated cortical neurons or *GBA1*-N370S dopaminergic neurons expressing mVenus-CAAX (Extended Data Fig. 4k-m). This data supports the use of tubulin as a marker of ectosomal buds by imaging. We then quantified the number of tubulin-positive ectosomal buds along the neurites in the PD mutant neurons; *GBA1* (N370S, L444P and W378G) and *LRRK2* (G2019S and R1441H). Increased ectosomal buds are present in all PD neurons assessed compared to their isogenic controls (Fig. 3g-k). Altogether, this data reveals ectosome shedding as a common feature of PD neurons with decreased GCase activity.

### Enhancing GCase activity mitigates ectosome formation in iPSC-derived neurons with *GBA1* and *LRRK2* mutations

PD neurons exhibit a negative correlation between GCase activity and the number of tubulin-positive ectosomes (Fig. 4a). This led us to explore the importance of GCase activity in ectosome formation. To this end, GCase activity was rescued in dopaminergic neurons using two approaches. First, PD neurons were treated with the GCase modulator 758 to restore GCase activity in *GBA1*-N370S and *LRRK2*-G2019S dopaminergic neurons (Fig. 4b-c) as previously reported^43,44^. Treatment with 758 effectively reduced the number of ectosomes shedding from PD neurons (Fig. 4d-f), with no effect on cell toxicity (Extended Data Fig. 5a-b). As a second strategy, wild-type human GCase (hGCase) was expressed in PD neurons along with an eGFP reporter under a separate promoter (Extended Data Fig. 5c-h). eGFP alone was expressed as a control (Extended Data Fig. 5c-h). Approximately 70-80% of iPSC-derived neurons were transfected as indicated by GFP fluorescence (Extended Data Fig. 5i-j) with no effect on cytotoxicity (Extended Data Fig. 5k-l). As expected, expression of hGCase restored GCase activity in both *GBA1*-N370S and *LRRK2*-G2019S dopaminergic neurons (Fig. 4g-h). Notably, expression of hGCase reduced the amount of ectosomes shedding from PD neurons back to that of their corresponding isogenic controls (Fig. 4i-k). These findings emphasize ectosome shedding as a direct consequence of reduced GCase activity.

**Fig. 4.**
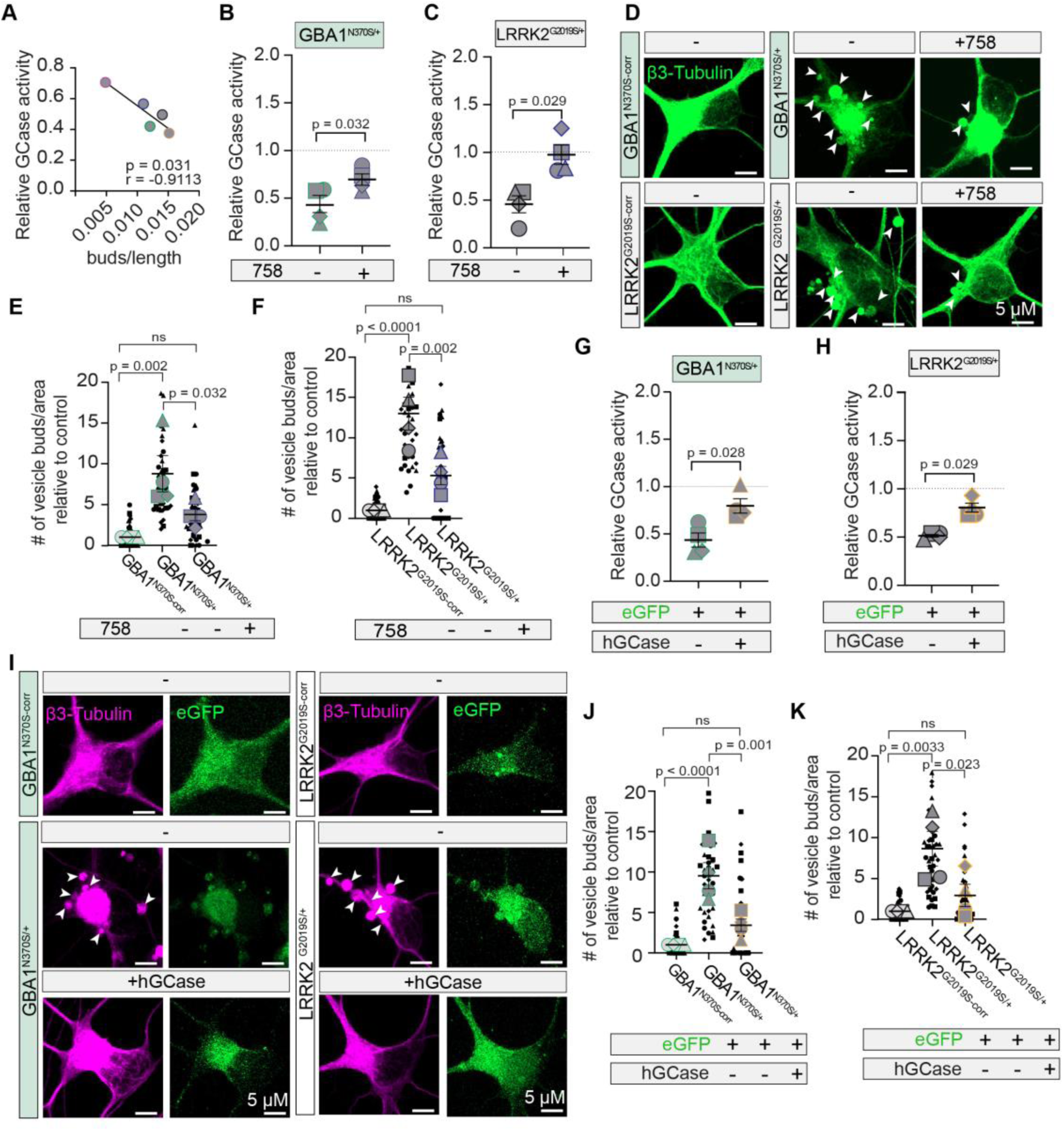
Increasing GCase activity suppresses ectosome formation from PD neurons. **(A)** Pearson correlation between GCase activity and the number of tubulin-positive ectosome buds in dopaminergic neurons with mutations in *GBA1* (N370S, L444P and W378G) and *LRRK2* (G2019S and R1441H). Dots represent the average reduced GCase activity and lengths per bud per line of 4 independent inductions. **(B-C)** GCase activity of *GBA1*-N370S and *LRRK2-*G2019S dopaminergic neurons with GCase modulator 758. 4 independent inductions; mean ± SEM; Mann-Whitney test. **(D)** Maximum intensity projections of *GBA1*-N370S and *LRRK2-*G2019S dopaminergic neurons and their isogenic controls +/− 758 treatment, stained for β3-tubulin. Arrows indicate ectosomes budding from the neurite and soma. (**E-F**) Quantification of tubulin-positive buds relative to control. *n* = 4 independent experiments; 10 cells per experiment; Mean ± SEM; One-way ANOVA with Šidák’s multiple comparison. Graph is depicted as a superplot where biological replicates are shown in large shapes, and technical replicates are shown as small shapes. **(G-H)** GCase activity of *GBA1*-N370S and *LRRK2-*G2019S dopaminergic neurons expressing eGFP or eGFP/hGCase. 4 independent inductions; mean ± SEM; Mann-Whitney test. **(I)** Maximum intensity projections of *GBA1*-N370S and *LRRK2-*G2019S dopaminergic neurons and their isogenic controls, expressing eGFP or eGFP/hGCase, stained for β3-tubulin. Arrows indicate tubulin-positive ectosomes budding from the neurite and soma. (**J-K**) Quantification of tubulin-positive buds relative to control. *n* = 4 independent experiments; 10 cells per experiment; Mean ± SEM; One-way ANOVA with Šidák’s multiple comparison. Graph is depicted as a superplot where biological replicates are shown in large shapes and technical replicates are shown as small shapes.

### Glucosylceramide-induced ectosomes are loaded with α-synuclein

A proposed characteristic of PD is the propagation of α-synuclein pathology. α-synuclein localizes to multiple regions within neurons allowing for several potential routes of release. For example, α-synuclein within lysosomes and multivesicular bodies could be released via lysosomal exocytosis and exosome secretion, respectively^45–47^. However, α-synuclein is also abundant in the cytoplasm where it interacts with plasma membrane sphingolipids associated with ectosome biogenesis^24–28^. This prompted us to explore whether α-synuclein is released within ectosomes.

Pathogenic fibrils, as opposed to soluble monomers, induce misfolding of endogenous α-synuclein^48,49^. We prepared and validated human α-synuclein preformed fibrils (PFFs) and confirmed their misfolding using a Thioflavin T assay. Here, an increase in the emission peak at 482 nm confirms amyloidogenic fibrils which are absent from monomeric α-synuclein (Extended Data Fig. 6a-b). Transmission electron microscopy was also performed to determine the length of labelled and unlabeled sonicated PFFs (Extended Data Fig. 6c-e). The majority of PFFs are shorter than 50 nm (Extended Data Fig. 6d-e). These shorter fibrils are more effective at seeding pathology compared to larger fibrils^50^. Post-translational modification of α-synuclein, most notably phosphorylation on residue Ser129, is associated with fibrillization and disease^51–53^. Rat cortical neurons were treated with PFFs and assessed for α-synuclein Ser129 phosphorylation by immunofluorescence. Both fluorescently tagged and unlabeled PFFs induced Ser129 phosphorylation which increased over time, while PBS or monomeric α-synuclein had no effect (Extended Data Fig. 6f-h).

Using live-cell super-resolution imaging, we discovered a massive accumulation of fluorescently tagged Atto594-PFFs within ectosomes budding from the plasma membrane of neurons treated with NBD-glucosylceramide (Fig. 5a and Extended Data Fig. 6i) and within vesicles released in the media (Fig. 5b). Atto594-PFFs can be seen close to the plasma membrane as they incorporate into growing ectosomes (Fig. 5c). This may promote enrichment of α-synuclein into ectosomes based on their affinity for plasma membrane sphingolipids^25–28^. The total amount of vesicles containing Atto594-PFFs with NBD-glucosylceramide treatment was approximately 30% (Fig. 5d). Even more ectosomes (60%) contained Atto594-PFFs when neurons were treated with the GCase inhibitor CBE compared to control neurons (Fig. 5e-f). This is likely due to the longer treatment time needed for CBE to induce vesicles which could increase the probability of α-synuclein interacting with the plasma membrane. The addition of PFFs to cortical neurons treated with CBE or NBD-GlcCer did not affect cytotoxicity (Extended Data Fig. 7a-b). Together, these data reveal glucosylceramide-induced ectosomes as a vehicle for α-synuclein release from neurons.

**Fig 5.**
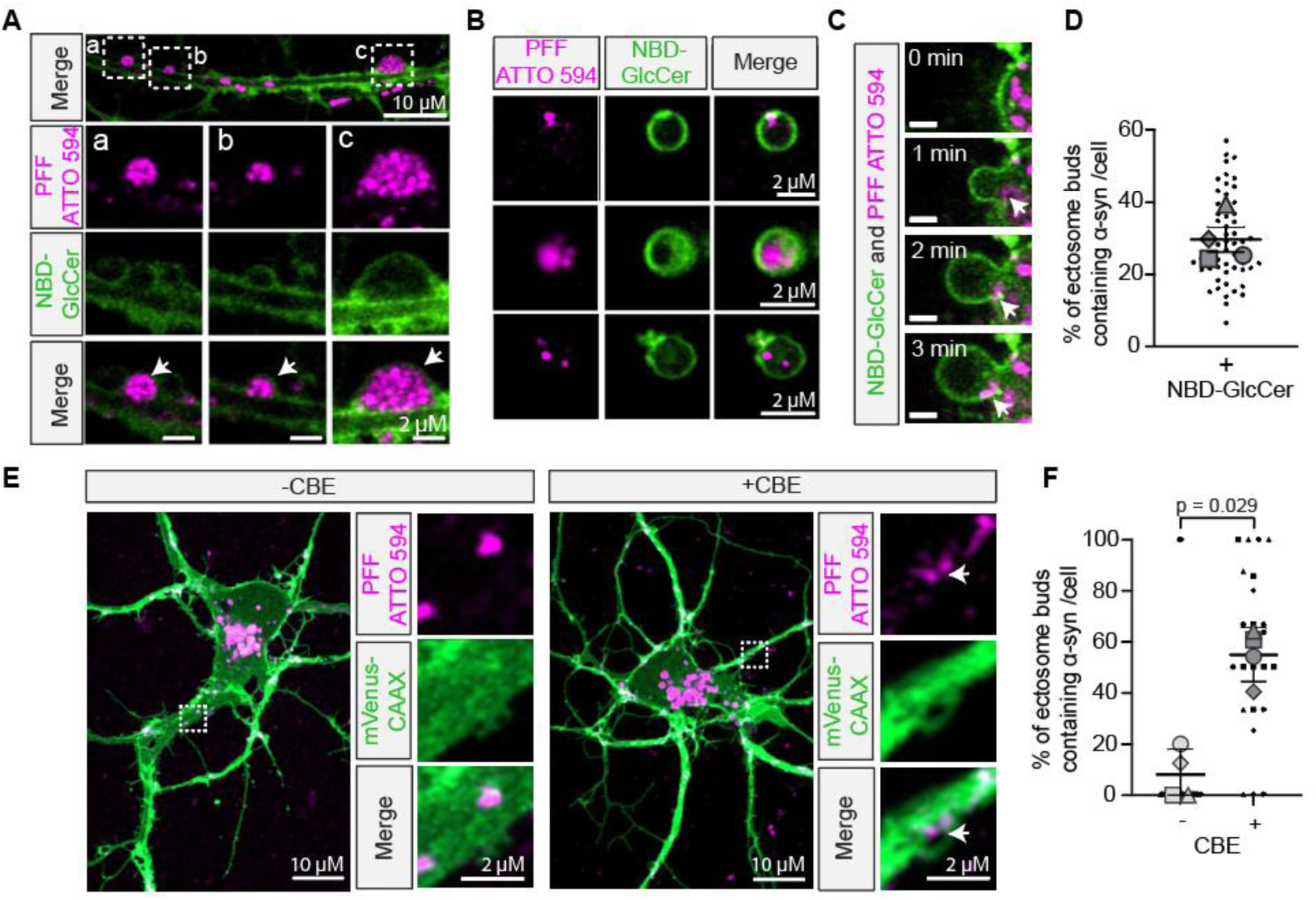
Ectosomes from cortical neurons accumulate α-synuclein fibrils. **(A)** Live-cell image of cortical neurite treated with NBD-GlcCer showing accumulation of ATTO594 α-synuclein fibrils within ectosomes and magnified images below. **(B)** Images of NBD-GlcCer-positive vesicles from neuronal conditioned media containing ATTO594 α-synuclein fibrils. **(C)** Time-lapse imaging showing ATTO594-PFF entering ectosome upon addition of NBD-GlcCer. **(D)** Percentage of ectosome buds filled with ATTO594-PFF. *n* = 4 independent experiments; 10 cells/experiment. Mean ± SEM. Graph is depicted as a superplot where biological replicates are shown in large shapes and technical replicates are shown as small shapes. **(E)** Live-cell image of ATTO594-PFF in ectosomes formed with CBE treatment in cortical neurons expressing mVenus-CAAX. Magnified images on the right. **(F)** Percentage of ectosomes filled with ATTO594-PFF with CBE treatment. *n* = 4 independent experiments; 10 cells/experiment. Mean ± SEM; Mann-Whitney test. Graph is depicted as a superplot where biological replicates are shown in large shapes and technical replicates are shown as small shapes.

### Ectosomes released from PD neurons are loaded with pathogenic α-synuclein

We next tested if ectosomes from PD neurons are similarly loaded with α-synuclein. Dopaminergic *GBA1*-N370S neurons and their isogenic controls expressing mVenus-CAAX were treated with α-synuclein PFFs and imaged live. *GBA1*-N370S neurons continue to form ∼5x more ectosomal buds compared to isogenic controls in the presence of PFFs (Fig. 6a-b). Of the ectosomes that form, more vesicles budding from *GBA1*-N370S neurons contain Atto594-PFFs compared to their isogenic controls (Fig. 6c). Large vesicles containing Atto594-PFFs were also detected in the dopaminergic neuron-conditioned media, indicating they are released (Fig. 6d). Interestingly, addition of Atto594-PFFs was sufficient to increase ectosomal buds in the dopaminergic control neurons (Extended Data Fig. 7c). Unlike rat neurons which were resistant to toxicity from PFF treatment (Extended Data Fig. 7a-b), human *GBA1*-N370S neurons and isogenic controls have a slight increase in cytotoxicity upon PFF treatment (Extended Data Fig. 7d). Altogether, these data confirm that PD neurons release ectosomes loaded with α-synuclein.

**Fig. 6.**
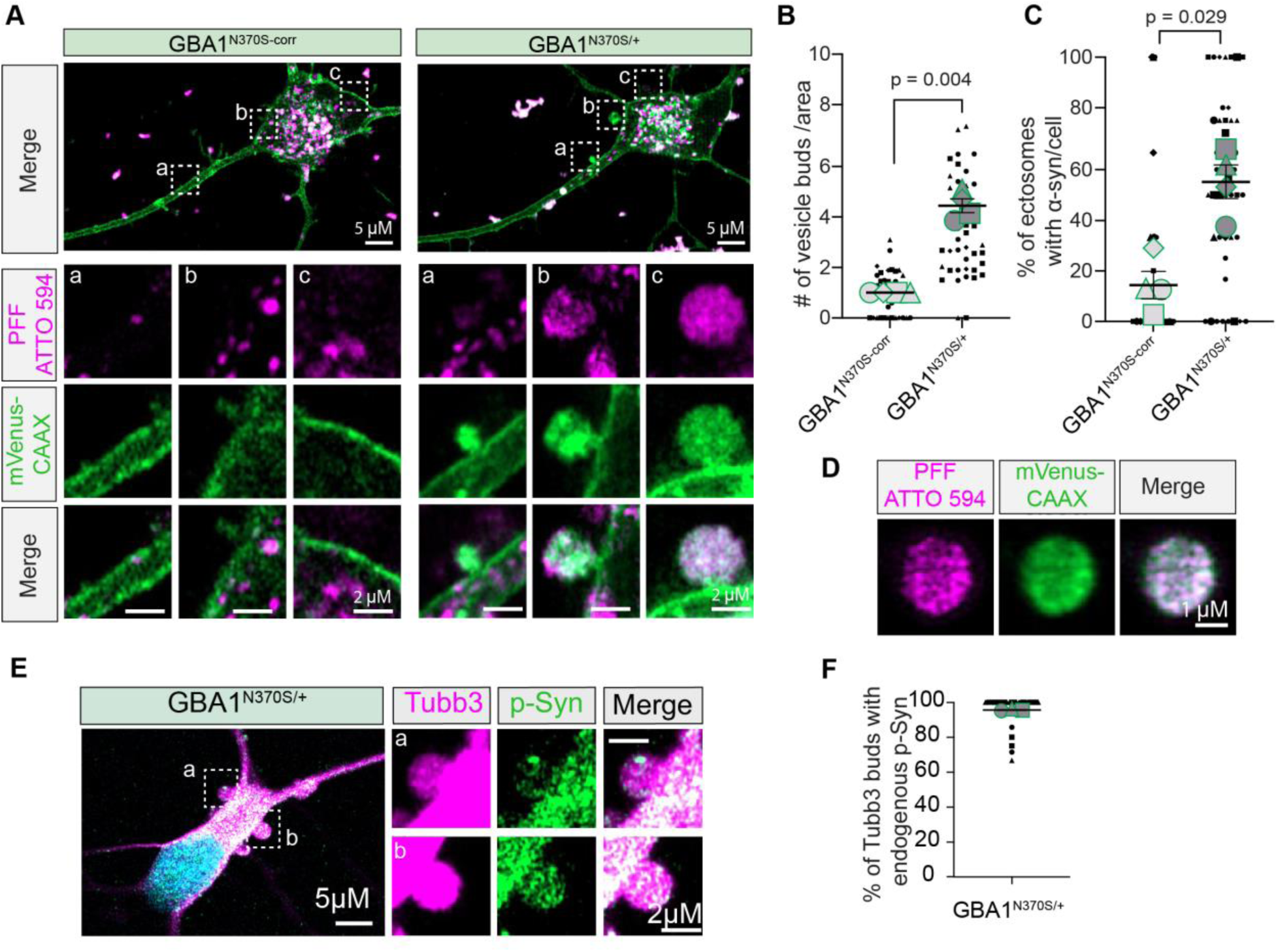
Ectosomes shed by PD neurons contain pathogenic α-synuclein. **(A)** Live-cell image of ATTO594-PFF in ectosomes from *GBA1*-N370S dopaminergic neurons but not isogenic corrected (corr) controls. Boxed areas are magnified below. **(B)** Number of mVenus-CAAX positive vesicles per area in *GBA1*-N370S relative to control neurons. *n* = 4 independent experiments; 10 cells/coverslip/treatment; mean ± SEM; One sample t-test. Graph is depicted as a superplot where biological replicates are shown in large shapes and technical replicates are shown as small shapes. **(C)** Percentage of ectosomes containing ATTO594-PFF in *GBA1*-N370S and control neurons. *n* = 4 independent experiments; 10 cells/experiment. Mean ± SEM; Mann-Whitney test. **(D)** Images of ATTO594-PFF containing vesicle from conditioned media of *GBA1*-N370S neurons. **(E)** Image of fixed *GBA1*-N370S dopaminergic showing endogenous pS129-α-synuclein in tubulin-positive ectosomal bud. Boxed areas are magnified on the right. **(F)** Percentage of tubulin-positive buds containing endogenous pS129-α-synuclein in *GBA1*-N370S dopaminergic. *n* = 4 independent experiments; 10 cells/experiment. Mean ± SEM. Graph is depicted as a superplot where biological replicates are shown in large shapes and technical replicates are shown as small shapes.

Since phosphorylation on Ser129 is associated with pathogenic fibrillization and disease^51–53^, we wondered whether α-synuclein within ectosomes is phosphorylated. We first turned to differential ultracentrifugation^23^ to test the abundance of α-synuclein in small versus large EVs. Total α-synuclein was detected in all fractions including large vesicles, small vesicles and the supernatant following high speed centrifugation likely corresponding to low molecular weight α-synuclein (Extended Data Fig. 7e-g). Next, we looked at α-synuclein phosphorylated on Ser129. *GBA1*-N370S neurons produced and released more phosphorylated α-synuclein compared to their isogenic controls (Extended Data Fig. 7f). This was expected as PD mutations in *GBA1* promote α-synuclein modification and/or fibrillization^29^. Unlike total α-synuclein, phosphorylated α-synuclein is enriched in the large vesicle fractions with little to no detectable signal in small vesicles or the supernatant (Extended Data Fig. 7f). Based on the size distributions of ectosomes determined above, we reasoned that ectosomes would be enriched in these large vesicle fractions. To confirm this, we imaged ectosomal buds from *GBA1*-N370S neurons treated with PFFs and indeed observed phosphorylated α-synuclein (Fig. 7h-i). Even in the absence of PFFs, *GBA1*-N370S neurons incorporate endogenous phosphorylated α-synuclein into ectosomes (Fig. 6e-f). Collectively, these data show that ectosomes released by PD neurons carry pathogenic α-synuclein.

**Fig. 7.**
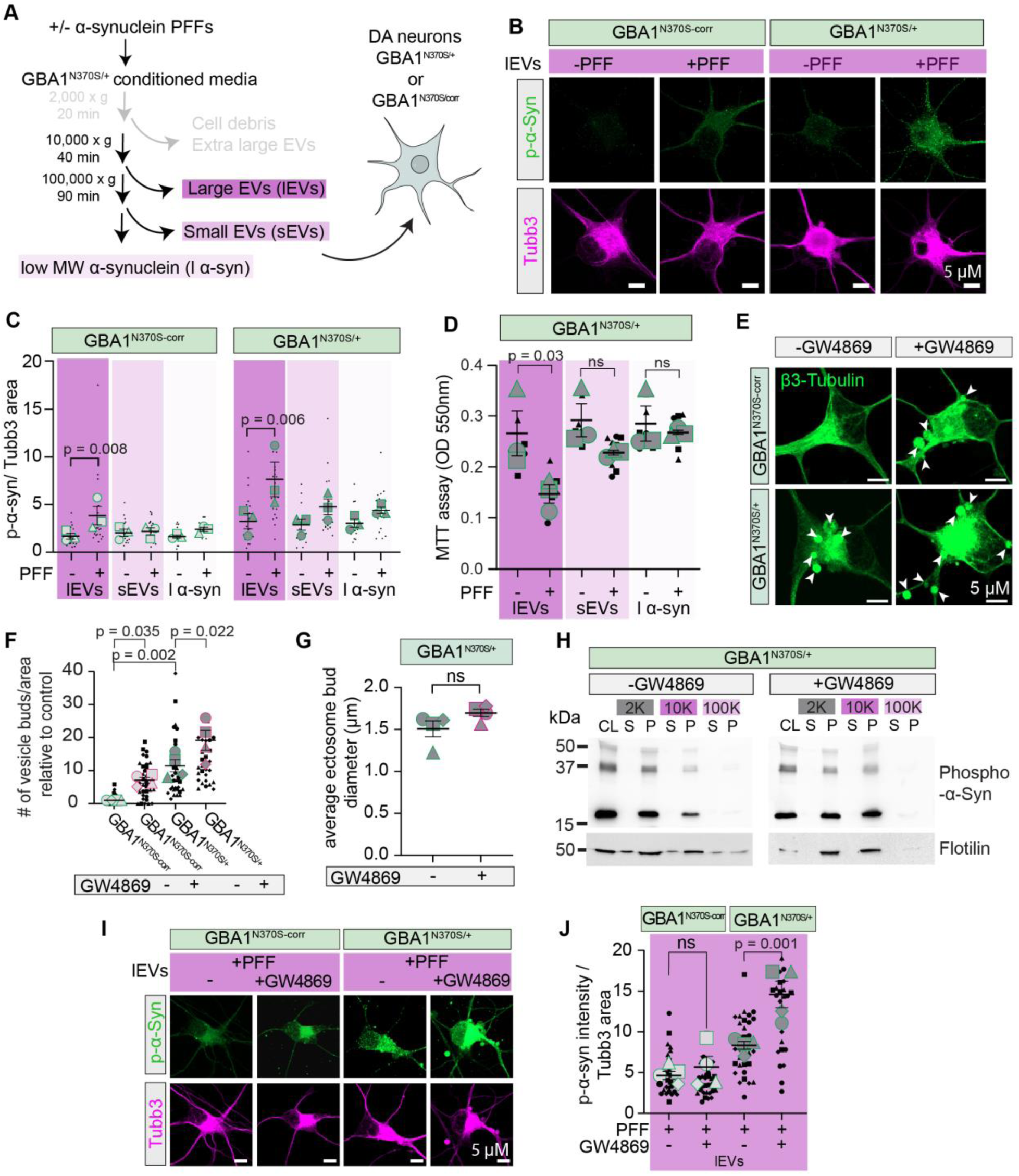
Ectosomes shed by PD neurons transmit α-synuclein pathology. **(A)** Schematic of assay to test effects of vesicles released by *GBA1*-N370S neurons. **(B)** Maximum intensity projections of *GBA1*-N370S and control neurons incubated with large vesicle fraction for 7 days derived from *GBA1*-N370S +/− PFF and stained with β3-tubulin and pS129-α-synuclein. **(C)** Quantification of the amount of pS129-α-synuclein normalized to the neuron area in *GBA1*-N370S and control neurons treated with different α-synuclein containing fractions for 7 days. *n* = 3 independent experiments; 6 cells/experiment. Mean ± SEM; One-way ANOVA with uncorrected Fisher LSD. Graph is depicted as a superplot where biological replicates are shown in large shapes and technical replicates are shown as small shapes. **(D)** Assessment of neuronal toxicity following treatment with large vesicles, small vesicles, or low molecular weight α-synuclein fractions released by PFF or PBS-treated *GBA1*-N370S neurons. *N* = 3 independent experiments; 3 technical replicates/experiment. Mean ± SEM; One-way ANOVA with Šidák’s multiple comparison. Graph is depicted as a superplot where biological replicates are shown in large shapes and technical replicates are shown as small shapes. **(E)** Maximum intensity projections of fixed *GBA1-*N370S dopaminergic neurons and its isogenic control +/− GW4869, stained for β3-tubulin. Arrows indicate tubulin-positive ectosomes budding from the neurite and soma. **(F)** Quantification of tubulin-positive buds relative to control. *n* = 4 independent experiments; 10 cells per experiment; Mean ± SEM; One-way ANOVA with Šidák’s multiple comparison. Graph is depicted as a superplot where biological replicates are shown in large shapes and technical replicates are shown as small shapes. **(G)** Average size of vesicle diameter of *GBA1-*N370S +/− GW4869. *n* = 4 independent experiments; bars show mean ± SEM; Mann-Whitney test. **(H)** Western blot of centrifugation fractions derived from GBA1-N370S neurons treated with PFF +/− GW4869 and immunoblotted for flotilin-1 and pS129-α-synuclein. CL, cell lysate; S, Supernatant; P, Pellet. 2K, 2000 x g; 10K, 10,000 x g; 100k, 100,000 x g. n=3 independent experiments. **(I)** Maximum intensity projections of GBA1-N370S and control neurons incubated with large vesicle derived from GBA1-N370S +/− GW4869 and PFFs for 7 days and stained with β3-tubulin and pS129-α-synuclein. **(J)** Quantification of the amount of pS129-α-synuclein normalized to the neuron area in GBA1-N370S and control neurons treated with large vesicle fraction +/− GW4869. n = 4 independent experiments; 8 cells/experiment. Mean ± SEM; One-way ANOVA with uncorrected Fisher’s LSD. Graph is depicted as a superplot where biological replicates are shown in large shapes and technical replicates are shown as small shapes.

### Ectosomes facilitate the transmission of pathogenic α-synuclein from PD neurons

Since ectosomes contain pathogenic α-synuclein fibrils, they could transmit α-synuclein pathology to other neurons. To test this, we incubated neurons with large vesicles, small vesicles, or the supernatant containing low molecular weight α-synuclein released from PFF or PBS-treated *GBA1*-N370S neurons (Fig. 7a). Large vesicles released by PFF-treated *GBA1*-N370S neurons increased α-synuclein phosphorylation in recipient *GBA1*-N370S after 7 and 14 days of incubation (Fig. 7b-c and Extended Data Fig. 8a-b). Small vesicles, on the other hand, only increased α-synuclein phosphorylation in the *GBA1*-N370S neurons after 14 days (Extended Data Fig. 8b), while low molecular weight α-synuclein had no effect (Fig. 7c and Extended Data Fig. 8b). Consistent with the propagation of α-synuclein pathology in the brain, large vesicles released by the recipient *GBA1*-N370S neurons can increase α-synuclein pathology in a third batch of neurons (Extended Data Fig. 8c-e). Finally, we assessed whether ectosomes released from PFF treated *GBA1*-N370S neurons induce neuronal dysfunction in recipient cells. Indeed, large vesicles, but not small vesicles or low molecular weight α-synuclein, released from PFF-treated *GBA1*-N370S increased neuronal toxicity in recipient neurons (Fig. 7d). These results indicate that large vesicles containing PFFs propagate α-synuclein and neuronal dysfunction.

Since the identity of vesicles cannot be determined based on size alone, it is possible that additional EV subtypes could be present in the large vesicle fraction. Exosomes for example, though typically smaller, have also been proposed to participate in the propagation of α-synuclein. To directly compare the contribution of ectosomes and exosomes, we took advantage of a widely used neutral sphingomyelinase inhibitor GW4869 that prevents exosome formation^54,55^. In addition to reducing exosomes, GW4869 also stimulates ectosome shedding^56,57^. We confirmed that GW4869 similarly increases the number of ectosomal buds in *GBA1*-N370S dopaminergic neurons and isogenic control neurons (Fig. 7e-f). GW4869 does not affect the size of ectosomal buds which are similar to those described above, (Fig. 7g and Extended Data Fig. 8f-h), nor does it affect cell toxicity (Extended Data Fig. 8i). We found that GW4869 increased the amount of phosphorylated α-synuclein in the large vesicle fraction consistent with more ectosomes, while decreasing the amount of flotillin in the small vesicle fraction consistent with less exosomes (Fig. 7h). Finally, we incubated neurons with large vesicles collected from PFF-positive *GBA1*-N370S treated with or without GW4869. We found a significant increase in phosphorylated α-synuclein after receiving large vesicles derived from GW4869-treated neurons (Fig. 7i-j). This data strongly supports ectosomes as the predominant carrier for propagating α-synuclein pathology between PD neurons.

## DISCUSSION

Extracellular vesicle formation is linked to changes in lipid metabolism and may play a crucial role in PD^30,58,59^. However, distinguishing the different subtypes of vesicles has posed a major challenge to the field^23^. Here, we employed a live-cell imaging approach to directly monitor ectosome formation in real time. We discovered that ectosomes are the predominant vehicle in transmitting pathogenic α-synuclein in cellular models of PD.

Other types of extracellular vesicles have been proposed to play a role in this process. Exosomes, for example, form via the fusion of multivesicular bodies with the plasma membrane, thereby releasing intraluminal vesicles into the extracellular space. Exosomes have been extensively studied in health and disease, including the release of pathogenic proteins such as α-synuclein^47^. They also mediate the transport of glucosylceramide from neurons to glia for degradation^3,30^. Many of these studies, however, are based on the use of exosome markers that have been identified on additional populations of extracellular vesicles. Perhaps the most notable example is CD63. In addition to its enrichment on exosomes, CD63 also localizes to the plasma membrane and can be released on plasma membrane derived ectosomes^60,61^. Here, we show that inhibiting exosomes promotes ectosome shedding and enhances the release of α-synuclein and transmission of α-synuclein pathology. This points to ectosomes as the predominant vehicle for transmitting pathogenic proteins and cautions against the use of exosome inhibitors alone as a therapeutic strategy, as this could exacerbate pathology^62^.

While we show that elevated glucosylceramide, due to reduced GCase activity, is sufficient to trigger ectosome formation, it remains possible that additional sphingolipids could be involved. For example, GCase also catalyzes the conversion of glucosylsphingosine to ceramide, such that PD mutations in *GBA1* could similarly increase glucosylsphingosine^10,16,29^. Alternatively, glucosylceramide can be converted to glucosylsphingosine by ceramidase^3^. Like glucosylceramide, glucosylsphingosine is also localized on plasma membrane rafts^63,64^ and elevated levels promote α-synuclein pathology in PD with mutations in *GBA1*^16,29,65^. This could indicate a potential role for this lipid in ectosome formation as well.

A notable feature of PD is the aggregation and potential prion-like spread of α-synuclein^66,67^. We show that this transmission of α-synuclein is due to ectosomes. But what remains to be determined is whether α-synuclein fibrils are recruited into growing ectosomes due to the altered lipid composition of the plasma membrane and/or specific protein machinery. Additionally, we observed α-synuclein fibrils is sufficient to induce ectosomal buds. This is consistent with the ability of pathogenic forms of α-synuclein to reduce GCase activity^72^. Alternatively, α-synuclein fibril interactions with the plasma membrane could affect local protein function and membrane curvature. It would be interesting to investigate how different α-synuclein strains affect the induction of ectosomes^68^. Conversely, glucosylceramide interacts with and promotes misfolding of cytosolic α-synuclein^13^. This could facilitate their incorporation into growing ectosomes. While endosomes/lysosomes containing α-synuclein fibrils could enter ectosomes as intraluminal vesicles. However, this only constituted a minority of the ectosomes observed, suggesting α-synuclein is more likely entering directly from the cytosol.

To summarize, this work reveals ectosomes as an important vehicle for the transmission of PD pathology. This has broad implications as similar sphingolipid alterations are found in additional PD backgrounds such as those with mutations in *SCNA*^11,12,69^, *PLA2G6*^70^, *PINK1*^71^, *VSP35*^70^ and even sporadic PD^19^. Given that GCase activity modulates neuronal susceptibility to α-synuclein pathology regardless of neuronal cell type, we propose that ectosome formation may be a common mechanism across different neuronal populations^72^. Ectosomes could even serve as a vehicle for the propagation of other toxic proteins, such as tau^73^. Therefore, lipid-induced ectosome formation likely contributes to the disease progression in PD and other related disorders.

## METHODS

### Animals

Experiments involving rat cells were approved by and performed under the Canadian Council of Animal Care at the University of Alberta (AUP#3358). Sprague-Dawley-timed pregnant rats were obtained from Charles River Laboratories and arrived at our facility 1 week before birth. Mouse experiments were approved by and performed under the Canadian Council of Animal Care at the University of Alberta (AUP#2711). Mice were group-housed in a temperature-controlled environment on a reverse 12-hour light-dark cycle and provided with ad-libitum food and water.

### Primary culture of cortical neurons

Mixed sex primary cortical cultures were prepared as previously described for hippocampal cultures^74^. Briefly, cortices were dissected from P0-P1 Sprague-Dawley rat pups and digested with papain (Fisher Scientific #LK003178) and Benzonase^®^ Nuclease (Sigma #E1014). After digestion, the tissue was gently triturated and filtered with a 70-µm nylon cell strainer (Fisher Scientific #22363548). Neurons were grown on poly-D-lysine (Sigma-Aldrich, #P6407) coated glass coverslips or plastic tissue culture dishes in neurobasal medium (Fisher Scientific, #21103049) containing B-27 supplement (Fisher Scientific #17504044), 2 mM GlutaMax (Fisher Scientific #35050061) and antibiotic-antimycotic (Fisher Scientific #15240062). 2 μM AraC was added to the media 2 days after plating to prevent the contamination of glial cells. Neurons were grown at 37°C in 5% CO_2_ and used at DIV 7 and DIV14.

### iPSC-derived dopaminergic neurons

Reprogramming of human iPSCs was completed by the Neuro’s EDDU, with PBMCs obtained from the Neuro’s C-Big Repository. The use of human iPSCs in the EDDU was approved by the McGill University Research Ethics Board (IRB Study Number A03-M19-22A). Human iPSC lines derived from PD patients with mutations in *GBA1* [N370S (female/65 yrs)^39^, W378G (female/ 62 yrs), and L444P (male/58 yrs)]^39^ and *LRRK2* [G2019S (female/62 yrs) and R1441H (male/54 yrs)] and their isogenic controls were obtained. Characterization and quality control data regarding the *GBA1* [L444P (male/58 yrs)] and *LRRK2* [G2019S (female/62 yrs) and R1441H (male/54 yrs)] iPSCs are provided in the Supplementary information. iPSCs were induced into dopaminergic neurons progenitors as described previously in^39,75^. CRISPR editing of patient-derived iPSCs to generate iPSCs was performed according to earlier published workflows^39,40^. Dopaminergic neural progenitor cells were dissociated with StemPro Accutase Cell Dissociation Reagent (ThermoFisher #A1110501) into single-cell suspensions. Cells were plated onto laminin (Fisher Scientific #23017015) and poly-L-ornithine (Sigma #P3655) coated coverslips in 6-well plates (1×10^5^ cells) for immunostaining and live-cell imaging or on coated 10 cm dishes (1×10^6^ cells) for vesicle collection by differential ultracentrifugation, with neural progenitor plating medium (DMEM/F12 supplemented with N2, B27 supplement; Fisher Scientific #11320082, 17502048, 17504044). Progenitors were differentiated into dopaminergic neurons by switching to differentiation medium (Fisher Scientific #21103049) supplemented with N2 and B27, BDNF (20 ng/mL; Peprotech #450-02), GDNF (20 ng/mL; Peprotech #450-10), Compound E (0.1 μM; Stemcell Technologies #73954), db-cAMP (0.5 mM; Cedarlane # ND07996), Ascorbic acid (200 μM; Sigma #A5960), TGF-β3 (10 ng/mL; Peprotech #100-63E) and laminin (1 μg/mL; Thermofisher #23017015). Mitomycin C (0.1μg/mL; Fisher Scientific, #AAJ63193MA) was added for 7 days to prevent glial contamination.

### Lipofectamine transfection of primary cortical neurons and dopaminergic neurons

Primary cortical neurons were transfected at DIV3. Dopaminergic neurons were transfected on the 3rd day of their final differentiation. Lipofectamine/DNA complexes were prepared according to the manufacturer’s instructions. Briefly, complexes were prepared in Opti-Mem (Gibco, #11058021) by mixing for each well of 6-well plates 1 μg of mCh-GPI plasmid (a gift from Dr. Aubrey Weigel, Janelia Research Campus, Virginia, USA^76^ or mVenus-CAAX plasmid (Addgene, #60411) or hGCase (Vectorbuilder, pRP[Exp]-EGFP-EF1A>hGBA1 [NM _0010 05741.3], Vector ID#VB900162) or GFP (pRRLsinPPTeGFP - gifts from Dr. Peter McPherson, McGill University, Canada) with 4.5 μl Lipofectamine3000 reagent (Fisher Scientific # L3000008). DNA and lipofectamine were each diluted with 125μl of Opti-Mem. Complexes were left for 15 min at room temperature, added to the cells for 3h followed by a complete media change.

### Lentivirus production and transduction

Lentiviral plasmids, pRRLsinPPTeGFP, pRSV-Rev, pMD2.g and pMDLg/pRRE were gifts from Dr. Peter McPherson, McGill University, Canada. Cloning mVenus-CAAX into pRRLsinPPT was performed using the Gibson NEBuilder^®^ Hifi DNA assembly kit. mVenus-CAAX was amplified with the following primer sequences: 5’ GCTGTTTTGACCTCCATAGAAGACACCGACTCTACTAGAGGATCCATGGTGAGCAAGGGCG 3’ and 5’ TCTTTCACAAATTTTGTAATCCAGAGGTTGATTGTCGAGCTCGAGCTACATAATTACACACTTTGTCTTTGACTTCTT 3’. As a two-fragment Gibson cloning reaction, the amplicon was inserted into pRRLsinPPTeGFP linearized by XhoI and BamHI. Sanger sequencing was performed to verify plasmid sequence (Plasmidsaurus). Viral particles were produced in HEK293T cells as previously described^77^. Briefly, HEK293T cells were transfected with lentiviral plasmids by calcium phosphate transfection. Media containing viral particles 24 h, 36 h and 48 h post-transfection were collected and filtered with a 0.45-μm filter to remove cell debris and concentrated by centrifugation at 7441 × *g* for 16 h at 4°C. Dopaminergic neurons were transduced with lentivirus (MOI 2) on the 3rd day of their final differentiation. The media was replaced with half-fresh half-conditioned dopaminergic cell media after 3h and the cells were used on the 7^th^ or 14^th^ day of their final differentiation.

### CBE, NBD-GlcCer, SiR-tubulin, GCase modulator 758 and GW4869 administration

CBE: Conduritol-β-epoxide (CBE, Cayman #15216, MedChem Express #HY-100944) was reconstituted at 100 mM in dimethyl sulfoxide (DMSO) and stored at −20°C. Primary cortical neurons were treated overnight at DIV6 or DIV13 (when incubated with PFFs) prior to use. NBD-GlcCer: C6-NBD-GlcCer (CedarLane #23209-1) was dissolved in cold HBSS by vortexing prior to uptake. On DIV7 or DIV14 (when incubated with PFFs), cells were washed with cold Hank’s balanced salt solution (HBSS; Fisher Scientific #SH30226801) and incubated with 2 μM NBD-GlcCer at 37°C for 15 min, then washed 3x with warm HBSS (37°C). SiR-Tubulin: Cortical neurons (DIV7) or dopaminergic neurons (7 days post final differentiation), were incubated with 0.5 μM SiR-tubulin (Cedarlane #CY-SC002) in 2 mL neuron-conditioned media for 30 min at 37°C. GCase modulator 758 NCGC0018875 (758, Millipore Sigma #531660) was reconstituted at 10 mM in DMSO and stored stored at −20°C. Dopaminergic cells were treated every other day with 10 μM 758, starting on the 4^th^ day for 7 days. Specificity and effective target engagement of NCGC0018875 have been verified and published previously^43,44,78^. GW4869: a non-competitive inhibitor of neutral sphingomyelinase (Millipore Sigma #567715) was reconstituted at 360 mM in DMSO and stored stored at −20°C. Dopaminergic cells were treated every other day with 2 μM GW4869 by addition to the culture media, starting on the 4^th^ day for 7 days. Specificity of GW4869 have been verified and published previously^54,55^.

### α-synuclein fibril preparation and characterization

ATTO 594 α-synuclein PFFs (Stressmarq #SPR-322E-A594) and untagged α-synuclein PFFs were generated according to The Michael J. FOX Foundation protocol. Briefly, human α-synuclein monomers (Proteos #RP-003) were incubated at 37°C at 5 mg/mL in PBS with constant agitation (1000 rpm) for 7 days using a Thermomixer. This was followed by aliquoting and storage at −80°C until use. Prior to use, PFFs were diluted in PBS and sonicated (30s, 0.5s on/off at 30% power) using a UP50H ultrasonic processor. Negative stain electron microscopy: 100 μg/mL PFFs before and after sonication were added to glow discharged 200 copper mesh carbon grid, washed with ddH_2_O and stained with 3% uranyl acetate (Electron Microscopy Sciences #22400) for 1 min. PFFs were visualized using a 200 kv JEOL 2100 Transmission Electron Microscope and seed length was determined with Fiji-ImageJ2 and the skeleton plugin. Thioflavin T assay: In a 96-well plate, 2.5 µL of the PFF or monomeric α-synuclein (5mg/mL) or PBS was added to 95 μL of the 25 µM Thioflavin T (Cedarlane #32553-10) per well. The plate was incubated at room temperature for 15 min and fluorescence was measured at ex385/em445 and ex450/em482 using Synergy H1 Multimode Microplate Reader. Treat neurons: PFFs diluted to 2 μg/mL were added to cortical neurons (DIV7) or dopaminergic neurons 7 days post final differentiation, incubated for 24 h with PFFs and used at 4-10 days post-treatment as indicated.

### Immunostaining

Cells were fixed in warm 4% paraformaldehyde (PFA) for 25 min at room temperature, washed twice in PBS with 0.1% Triton X-100 and blocked with PBS containing 2% bovine serum albumin and 0.2% Triton X-100 for 1 h at room temperature. Cells were incubated with primary antibodies: anti-mouse anti-β3-tubulin (1:1000; Biolegend #801202), anti-chicken Tyrosine Hydroxylase (1:1000; EMD Millipore #AB9702) and anti-rabbit P-S129-synuclein (1:1000; Novus Biologicals #61121) diluted in blocking buffer overnight at 4°C. Cells were washed 3x in PBS with 0.1% Triton X-100 and incubated with secondary antibodies: goat anti-mouse Alexa Fluor 488 or 647 (1:1000; ThermoFisher #A-11001 and A-21235), goat anti-rabbit Alexa Fluor 488 (1:1000; ThermoFisher #A-11008) and goat anti-chicken Alexa Fluor 647 (1:1000; ThermoFisher #A-21449) diluted in blocking buffer for 1 h at room temperature. After washing, the cells were incubated with DAPI diluted in PBS for 10 min and mounted using DAKO fluorescence mounting media.

### GCase activity

Glucosylceramidase Activity Assay kit (Abcam #ab273339) was used as per manufacturer’s instructions. Fluorescence was measured (Ex/Em = 360/445nm) at 37°C in endpoint mode using Synergy H1 Multimode Microplate Reader (BioTek Instruments Inc.). Three technical replicates were performed for each experiment and all measurements were corrected by background subtraction. The activity of GCase in the sample was calculated from a 4-MU standard curve taking initial mg/mL protein concentration, sample volume and reaction time into account. Samples were normalized to control.

### Cell Death Assays

Neurons were plated on 96-well PDL-coated plates and treated O/N with 100 µM CBE or DMSO or for 15 minutes with 2 µM NBD-GlcCer or MeOH, followed by incubation in HBSS for 45 minutes, the maximum duration of our live-cell imaging. Ethidium Homodimer-1 Assay. Neurons were washed twice in warm PBS and incubated with 4 μM ethidium homodimer-1 (ThermoFisher #L3224) in PBS for 30 min at 37°C to label dead cells, washed twice in PBS and fluorescence was measured using Synergy H1 Multimode Microplate Reader (BioTek Instruments Inc.). Apoptosis assay: Cells were washed twice, fixed in 4% PFA, stained using a TUNEL assay kit (Abcam #ab66110) according to the manufacturer’s protocol, DAPI as a counterstain and imaged within 3 h of staining. TUNEL-positive cells were detected by thresholding images using the default algorithm and the number of particles was analyzed using Fiji-ImageJ software. For quantification, 2 tile scans (covering 465.5 x 465.5 μm) per coverslip were averaged.

### MTT assay

Cells (50,000 cells/well) were plated in 96-well flat bottom plates and subjected to CBE (Cayman #15216, MedChem Express #HY-100944), GW4869 (Calbiochem #567715), GCase modulator (NCGC0018875, Millipore Sigma #531660), exogenous C6-NBD-GlcCer (CedarLane #23209-1), SiR-tubulin (CedarLane #CY-SC002), untagged α-synuclein PFFs and/or a combination of those treatments mimicking our experimental setups. Subsequently, the MTT assay (Roche # 11465007001) was performed. Briefly, 10uL of 3-[4,5-dimethylthiazol-2-yl]-2,5-diphenyl tetrazolium bromide (MTT, 5mg/mL) in phenol red-free neurobasal media was added to each well. The plates were incubated for an additional 4 h at 37°C. After incubation, formazan crystals were solubilized in the provided solubilization buffer O/N at 37°C and the absorbance was measured at 550 nm using Synergy H1 Multimode Microplate Reader (BioTek Instruments Inc.).

### Separation of extracellular vesicles

The neuron-conditioned medium of dopaminergic neurons was collected on day 14 of the final differentiation. For blotting, protease and phosphatase inhibitors (Thermo Scientific Halt Protease Inhibitor Cocktail #78430) were added to the media. The media was then centrifuged at 2,000 × g at 4 °C for 20 min, the resulting supernatant was centrifuged at 10,000 × g at 4 °C for 40 min. The resulting supernatant was centrifuged at 100,000 × g at 4 °C for 90 min in a TLA-110 fixed angle rotor (k-factor 13, Beckman Coulter). Pellets were resuspended in ice-cold 25 µL lysis buffer (20 mM Tris pH 7.4, 100 mM NaCl, 5 mM EDTA, 1% Triton-X 100, supplemented with 1 × HALT protease and phosphatase inhibitor cocktail for Western blot. For treatment use, cells pellets resuspended in ice-cold PBS or in the final supernatant was given to dopaminergic neurons on the 3^rd^ day of their final differentiation and incubated for 7 or 14 days.

### Western blotting

Neurons on 10-cm plates were washed 3 times in PBS and lysed in lysis buffer (20 mM Tris pH 7.4, 100 mM NaCl, 5 mM EDTA, 1% Triton-X 100, supplemented with 1 × HALT protease and phosphatase inhibitor cocktail (ThermoFisher Scientific, #78430). Following centrifugation of EVs, pellets were resuspended in 25 µL lysis buffer and SDS-PAGE sample loading buffer, resolved by SDS-PAGE and processed for Western blotting. Primary antibodies used include rabbit polyclonal anti-syntenin (1:1000, Abcam #19903), mouse monoclonal anti-flotillin-1 (1:1000, BD Biosciences #610821), mouse monoclonal anti-β3-tubulin (1:1000, Biolegend #801202), rabbit monoclonal anti-CD81 (1:1000, Abcam #109201), rabbit monoclonal anti-Tomm20 (1:1000, Abcam # 186735), rabbit polyclonal anti-Calnexin (1:1000, Abcam # 227310), rabbit polyclonal anti-P-S129-synuclein (1:2000, Novus Biologicals #NBP2-61121), rabbit polyclonal anti-synuclein (1:2000, Novus Biologicals #NBP2-15365), mouse monoclonal anti-GCase (1:1000, Abcam #AB55080-1001), rabbit polyclonal anti-GFP (1:1000, ThermoFisher #A-6455) and mouse monoclonal anti-β-actin (1:5000, NEB #3700S). Prior to incubation with antibodies against synuclein and P-S129-synuclein, membranes were incubated in PBS containing 4% PFA and 0.1% glutaraldehyde for 30 min to increase sensitivity ^79^. HRP-conjugated affinity purified secondary anti-rabbit and anti-mouse were purchased from Jackson Immunoresearch Labs.

### Lipidomic analyses

At DIV6 cortical neurons were treated with DMSO or 100 µM CBE. On DIV7, cells were washed twice in ice-cold PBS, scraped into ice-cold 1:1 methanol:water, and stored in liquid nitrogen. Lipidomic analysis from 5 independent experiments per group by the Metabolomic Innovation Center. In brief, samples were spiked with NovaMT LipidRep Internal Standard Basic Mic for Tissue/Cells containing 15 deuterated lipids, dichloromethane, and methanol. Lipids were extracted using a modified Folch liquid-liquid extraction protocol, air dried, resuspended in NovaMT MixB and diluting in NovaMT MixA, LC-MS analyses were performed on the lipid extracts using Thermo Vanquish UHPLC linked to Bruker Impact II QTOF Mass Spectrometer. Data of samples were independently acquired in both positive and negative ionization modes with mass range m/z 150-1500. Data post-processing and normalization were performed using LipidScreener (Nova Medical testing Inc.).

### Mouse surgery

Ai9 male and female mice (Jackson Laboratory, Strain #:007909, RRID:IMSR_JAX:007909) were given 5 mg/kg of carprofen via ad-libitum water 24 hours before and for 72 hours after surgery. Mice were initially anesthetized in an induction chamber with 4% isoflurane mixed with pure oxygen and then maintained at 1.5-2.0% isoflurane when placed on the stereotaxic frame. An electric heating pad kept the body temperature at 37°C during the surgery. Mice were administered a local anesthetic (0.5% bupivacaine) subcutaneously before the skin was incised along the midline to expose the skull. The skull was cleaned and kept moist during surgery with 0.9% saline. The head was then levelled in rostral-caudal and medial-lateral directions according to bregma, allowing precise viral injection.

### Virus injection

Craniotomies were marked and manually drilled using a 400µm dental drill above the retrosplenial cortex. Coordinates were: Anterior-posterior −1.25 mm from bregma, mediolateral 0.7 mm from bregma, and −0.5 mm from brain surface. Injections were made by lowering pulled glass micropipettes (10 µm) backfilled with mineral oil and loaded with virus into the brain. 100 nl of AAVrg-hSyn-Cre (Addgene #105553, titer ≥ 7×10¹² vg/mL) was injected into the left hemisphere. After injection, the skin was sutured, and the mice returned to clean heated cages. This virus gets trafficked in the retrograde direction and drives expression of cre-recombinase in neurons in the contralateral hemisphere and results in td-tomato expression in Ai9 reporter mice^80^. Three weeks following virus injection, the skull of the mouse was thinned using a 1 mm dental drill bit. The skull was thinned until flexible, and the vasculature could be observed. A pulled glass micropipettes (10 µm) backfilled with mineral oil and loaded with NBD-GlcCer (4 μM/100 nl) and lowered into the right hemisphere near retrograde labelled, td-tomato expressing neurons.

### Two Photon Microscopy

Imaging was performed on an upright ThorLabs Bergamo II microscope with a Ti:Sapphire femtosecond laser (Thorlabs Tiberius) and 8 kHz resonant scanner (Thorlabs), with the laser set to 950 nm. Two-photon emission was filtered using a 562 nm long pass dichroic mirror (FF562-Di03-32×44, Semrock) and 525 nm emission filter (ff03-525/50-32, Semrock) before entering a GaAsP PMT (PMT2100, Thorlabs). The main microscope body was inside a light-tight enclosure with sound-dampening foam (5692T49, McMaster Carr) on all sides. Mice were placed in a half open plastic tube, with square white nesting pads to provide cushioning. A Nikon 16 x 0.8NA objective was used for 512 x 512px imaging, averaging 10 frames. Bidirectional correction was adjusted per field of view based on location and depth of imaging. Images were acquired at 17.1x zoom and z-stacks were taken with 1 µm spacing across a total of 16 stacks, resulting in a 53.34 x 53.34 x 16 µm field of view (0.10 x 0.10 x 1.0 µm/pixel). Laser power and PMT gain were set to 60-150mW and 10, respectively. Laser power was modulated through a quarter wave plate (Thorlabs Power Modulator). The mice were kept under anesthesia for the entire procedure by intraperitoneal injection with urethane (0.1g/mL) injection. The resulting imaging were binned (x/y shrink factor 2 and z shrink factor 1) and transformed by a 2-pixel Gaussian blur to reduce noise. Shifts in the Z-axis caused by motion during imaging were corrected by aligning a time-lapse of neurites across optical planes.

### Histology and tissue processing

Mice were anesthetized by intraperitoneal injection of urethane, intracardially perfused with ice-cold PBS, followed by 4% PFA in PBS. Brains were extracted and postfixed in 4% PFA for 24 hr and stored in PBS at 4°C. 50 μm coronal sections were obtained using a vibratome (Leica VT1000s, Germany). Slices were washed with PBS (3 × 15 min) and blocked using 2% bovine serum albumin in PBS-T (0.1% Triton X-100 in PBS) for 1 hr at room temperature. Sections were incubated with rabbit anti-NeuN (1:1000, Millipore #ABN78) in blocking buffer at 4°C overnight. Slices were washed with 0.1% PBS-T (3 x 15 min) and incubated with donkey anti-rabbit Alexa Fluor 647 (1:1000, ThermoFisher #A-31573) at room temperature for 4 hr in the dark. After washing with 0.1% PBS-T (3 × 10 min), incubation with DAPI (1:10,000, Abcam #ab2285429) in PBS for 30 min at room temperature and then PBS (2 × 10 min). Finally, slices were mounted using DAKO fluorescence mounting media.

### Super-resolution microscopy

Neurons expressing mCh-GPI, mVenus-CAAX and/or treated with CBE, NBD-GlcCer, SiR-tubulin or PFFs-ATTO-594 as described above were washed twice in warm PBS and imaged live in HBSS (Fisher Scientific #SH30226801). LSM900 with Airyscan2 equipped with a plan-apochromat 40x oil objective (Zeiss, NA = 1.3), incubation system to maintain 37°C, humidity and 5% CO_2_, and ZEN software (Zeiss). Images were acquired in the SR-4Y mode, frame scan using the Z-piezo stage. Timelapse movies were acquired at 12 frames/s. Time alignment was applied to correct for drift. 4 independent experiments and 10 cells per experiment were imaged. Vesicles from neurons treated as described above for 45 minutes were collected from neuron-conditioned media, allowed to settle on PDL-coated coverslip overnight at 4°C and imaged as above. One hundred vesicles derived from 4 independent experiments were imaged per condition. Neurons immunostained for β3-tubulin, tyrosine hydroxylase and P-S129-synuclein were imaged using a plan-apochromat 40x oil objective (Zeiss, NA = 1.3). Images related to the neurotransmission experiment were acquired using plan-apochromat 63× oil objective (Zeiss, NA = 1.4) in SR-2Y mode. 4 independent experiments and 10 cells per experiment were imaged. Coronal brain slices of mice were imaged with a plan-apochromat 63× oil objective (Zeiss, NA = 1.4) in the SR mode. Scan speed was set to 4 and frame time 34.53s. For larger overview images tile scanning was applied. Tiled images were stitched with 10% overlap. All images were Airyscan processed. Images related to percentage eGFP and hGCase/eGFP transfection in cells were acquired using plan-apochromat 20× air objective (Zeiss, NA = 0.8) in SR-4Y mode. 4 independent experiments and 4 tiles per coverslip (each covering an area of 2.21 x 2.21 mm) per experiment were imaged.

### In vitro image analysis

All images were analyzed using the Fiji-ImageJ2 (version 2.14.0) software. All analyses of vesicle buds were quantified blind. Movies were generated using Imaris AI Microscopy Image Analysis Software (Oxford instruments). Number of ectosomes at the soma and along the neurites were tracked through the different z-stacks and manually counted. The number of ectosomes was normalized against cell area as determined by thresholding (Huang algorithm) a maximum intensity projections of 3D image stacks. The number of ectosomes containing tdTomato, PFFs or SiR-tubulin were tracked through z-stacks, manually counted, and normalized against the total amount of ectosomes. Ectosomes with protein of interest at the neck only were counted as negative. The diameter of vesicles was measured in the optical section that bisected the middle of the vesicle. For attached ectosomes, the diameter of at least 15 vesicle buds per cell was determined. 10 images per experiment were taken and 4 independent experiments were conducted. The diameter of 100 released vesicles per experimental condition was determined. Phosphorylated α-synuclein was calculated as the percentage of cell area. The cell area was determined by thresholding by β3-tubulin (Huang algorithm), expanding that selection by 0.5 μM and clearing signal outside of the cell. The percentage of eGFP positive dopaminergic neurons, upon eGFP or hGCase expression, was determined by manually and normalized against the number of DAPI-positive nuclei per tile scan. 4 tile scans per experiment were taken and 4 independent experiments were conducted.

### In vivo image analysis

Maximum intensity projections of three-dimensional image stacks were created. Manual thresholding was applied to select soma and all neurites of the tdTomato expressing cell. The selection was enlarged by 2 μm and cleared. ‘Analyze particles’ was applied to determine the amount of tdTomato positive puncta larger than 0.5 μm. A 3D-quantification and colocalization of tdTomato- and NBD-GlcCer positive puncta within NeuN-positive cells was performed using the Fiji-ImageJ2 (version 2.14.0) software. Images were preprocessed by isotropic scaling setting voxel size to 0.1 μm in each dimension. Images were split into individual channels and duplicated. Next, a 3D Gaussian blur of 3.0 μm, 0.4 μm, and 0.4 μm was applied on the first copy of each NeuN, tdTomato, and NBD-GlcCer channel, respectively. On the second copy of each NeuN, tdTomato, and NBD-GlcCer channel, a 3D Gaussian blur of 1.5 μm, 0.4 μm, and 0.4 μm was applied respectively. The difference of Gaussians was taken by subtracting the second copy from the first for each channel. The contrast of each resulting channel was adjusted to 0.1 % of all pixels in the z-stack to be saturated. A mask for each channel was created using the MaxEntropy algorithm using the histogram of each z-stack. Then, 3D-masks were created for colocalization quantification. The AND image operation was performed between the mask of the NeuN and the mask of the NBD channels. This represents NBD-positive puncta contained within NeuN-positive cells. The OR image operation was performed between the mask of the tdTomato and the NBD channels. This represents tdTomato- and NBD-GlcCer double positive puncta. Finally, quantification of the 3D masks using the 3D Suite plugin was performed ^81^. Objects in the masks were segmented in 3D. NeuN, tdTomato and NBD-GlcCer double positive volumes and puncta counts were quantified. tdTomato- and NBD-GlcCer double positive puncta counts were confirmed manually and quantified per NeuN-positive cell based on the 3D masks.

### Statistical analyses

Statistical analysis and graphing were performed using GraphPad Prism 10. No data points were excluded from analyses. Researchers were blinded during analyses as indicated above. Data is presented as mean ± SEM, unless otherwise stated. Statistical analysis was performed on the experimental averages. To assess statistical significance, two-tailed unpaired t-test, Mann-Whitney test, one-way ANOVA with Dunn’s multiple correction test or uncorrected Fisher’s LSD were performed, as indicated in the figure legends.

### Reporting summary

Further information on research design is available in the Nature Portfolio Reporting Summary linked to this article.

## Supporting information

Supplementary information iPSC line validation

## Acknowledgments

We thank Dr. Peter McPherson, Dr. Hugo Bellen, Dr. Guang Lin, Dr. Xueyang Pan, Dr. Mingxue Gu, Dr. Michal Tyrlik, Dr. Ameya Walimbe, and Dr. Shree Kaduskar Dr. Sue-Ann Mok for helpful comments on the manuscript. We thank Dr. Xuejun Sun and Pinzhang Gao for assistance with EM sample preparation, Dr. Andrew Simmonds for the use of Imaris software. Kristiaan Jacquemyn for the support related to serialEM software. We thank C-BIG for facilitating access to the patients and patient-derived materials. Experiments were performed at the Faculty of Medicine & Dentistry Cell Imaging Core RRID: SCR_019200.

## Author contributions

Conceptualization: MSI, Julie Jacquemyn (JJ-1). Methodology: MSI, JJ-1, BM, Jesse Jackson (JJ-2), CXQC, TD. Investigation: JJ-1, CG, JW, NYL, LFRA, JC, KC, CAM. Resources: NA, MN, ED, ZY. Visualization: MSI, JJ-1. Funding acquisition: MSI, JJ-1, TD. Supervision: MSI, JJ-2, TD. Writing – Original Draft: MSI, JJ-1. Writing – Review & Editing: MSI, JJ-1, JJ-2, CXQC, TD.

## Supplementary Materials

Supplementary information - Characterization and Quality control of *GBA1* L444P and *LRRK2*-G2019S and *LRRK2*-R1441H iPSCs

Video S1 - Release of tdTomato-positive vesicles from neurons in vivo

Table S1 - List of significantly altered HexCer species

Table S2 - Statistical analysis sample

---

**Extended Data Figure 1.**
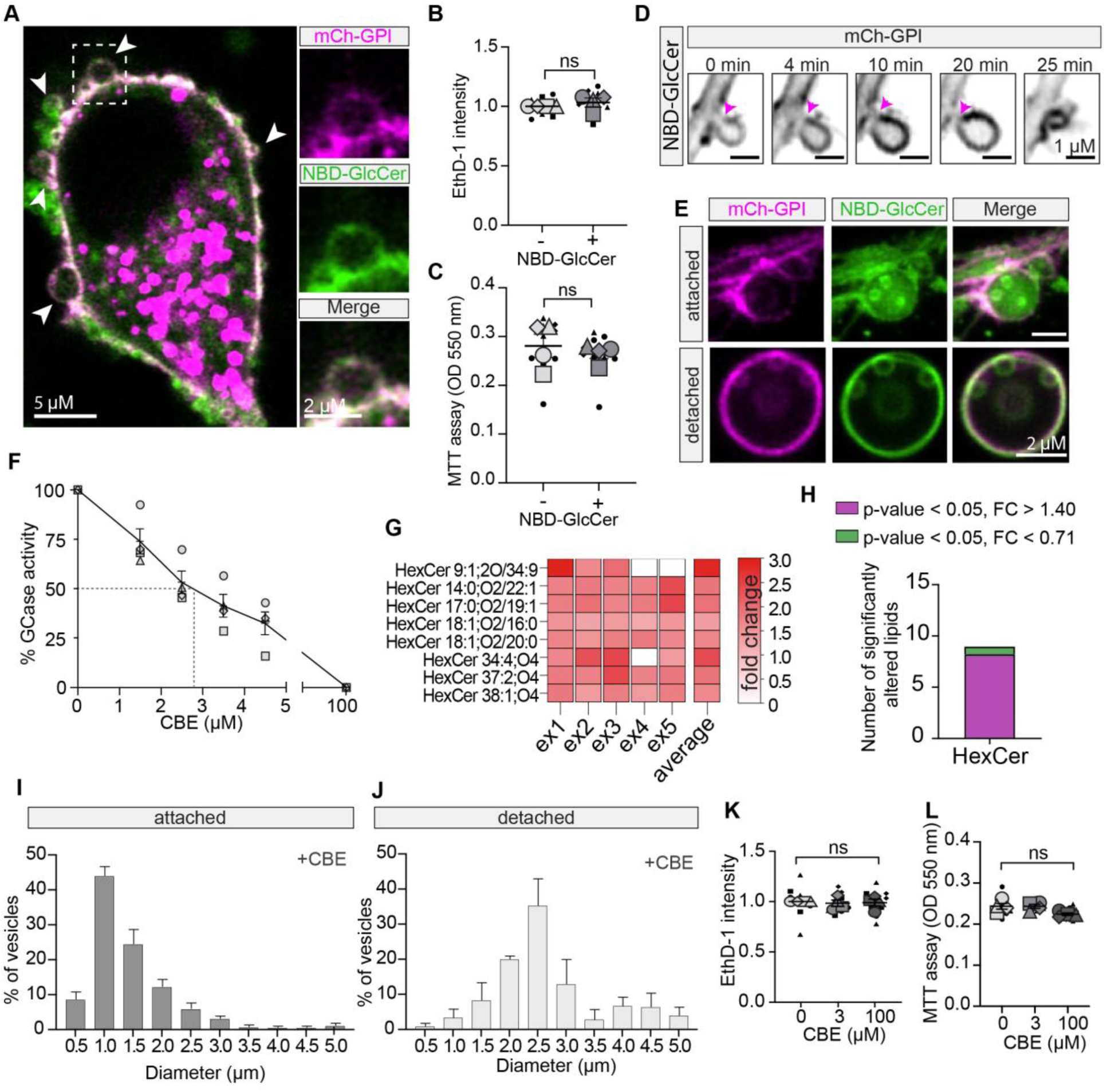
Increased GlcCer induces ectosome formation. **(A)** Live-cell image of cortical neuron expressing mCh-GPI treated with NBD-GlcCer. Boxed area highlighting vesicles budding from soma and magnified on the left. **(B)** Intensity of ethidium homodimer-1 (EthD-1) staining of neurons treated with NBD-GlcCer relative to control. *n* = 4 independent experiments; mean ± SEM; One sample t-test. Graph is depicted as a superplot where biological replicates are shown in large shapes and technical replicates are shown as small shapes. **(C)** Cytotoxicity assessment of NBD-GlcCer on cortical neurons by MTT assay. *n* = 4 independent experiments; 3 technical replicates/experiment; mean ± SEM; Mann-Whitney test. Graph is depicted as a superplot where biological replicates are shown in large shapes and technical replicates are shown as small shapes. **(D)** Time-lapse imaging showing mCh-GPI positive ectosome biogenesis with NBD-GlcCer treatment. **(E)** Images of attached and detached ectosomes containing intraluminal vesicles labelled with mCh-GPI and NBD-GlcCer from the neuron-conditioned media. **(F)** GCase activity of cortical neurons treated with CBE. The plotted line is a non-linear fit of values. *n* = 4 independent experiments. **(G)** Heatmap showing fold change of significantly altered HexCer (galactosylceramide and glucosylceramide) of neuron treated with 100 µM CBE. *n* = 5 independent experiments. **(H)** Average fold change of HexCer species with CBE treatment. *n* = 5 independent experiments. (**I-J**) Size distribution of attached and detached mCh-GPI-positive vesicles from neurons treated with 100 µM CBE. *n* = 4 independent experiments; mean ± SEM. (**K-L**) Neurons were treated with CBE and stained with ethidium homodimer-1 (EthD-1) and assessed for cytotoxicity by MTT assay. *n* = 4 independent experiments; mean ± SEM; One-way ANOVA with Dunn’s multiple comparisons test. Graph is depicted as a superplot where biological replicates are shown in large shapes and technical replicates are shown as small shapes.

**Extended Data Figure 2.**
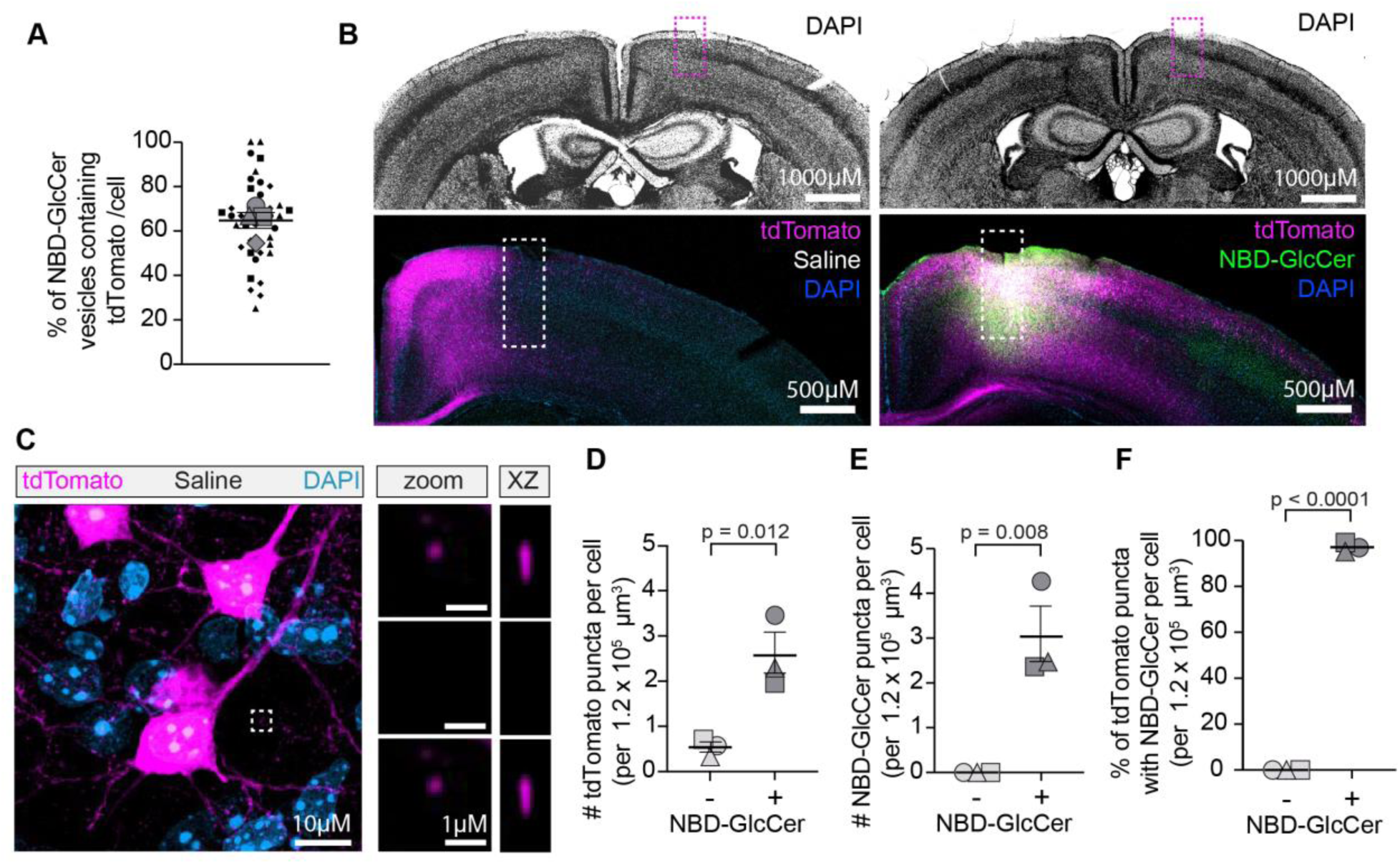
Ectosomes containing tdTomato fluorophore are released by cortical neurons treated with glucosylceramide in vivo. **(A)** Percentage of NBD-GlcCer positive buds containing the tdTomato fluorophore. *n* = 4 independent experiments; mean ± SEM. Graph is depicted as a superplot where biological replicates are shown in large shapes and technical replicates are shown as small shapes. **(B)** Coronal section of mouse brain with saline or NBD-GlcCer injection. Tiled widefield and fluorescent images are displayed. Boxes indicate the saline or NBD-GlcCer injection side. **(C)** Maximum intensity projection of fixed tissue showing a cortical neuron expressing tdTomato after saline administration. Boxed area shows magnification of tdTomato positive vesicle, ≥ 2 µM from neuron, with orthogonal views on the right confirming it is fully detached. **(D)** Number of tdTomato positive puncta in NeuN positive cells. *n* = 3 animals; 5 images per n; mean ± SEM; Unpaired t test.. **(E)** Number of NBD-GlcCer positive puncta in NeuN positive cells. *n* = 3 animals; 5 images per n; mean ± SEM; Unpaired t test. (**E**) Percentage of tdTomato positive puncta with NBD-GlcCer in Neun positive cells neighboring the tdTomato positive cortical neuron. *n* = 3 animals; 5 images per n; mean ± SEM; Unpaired t test.

**Extended Data Figure 3.**
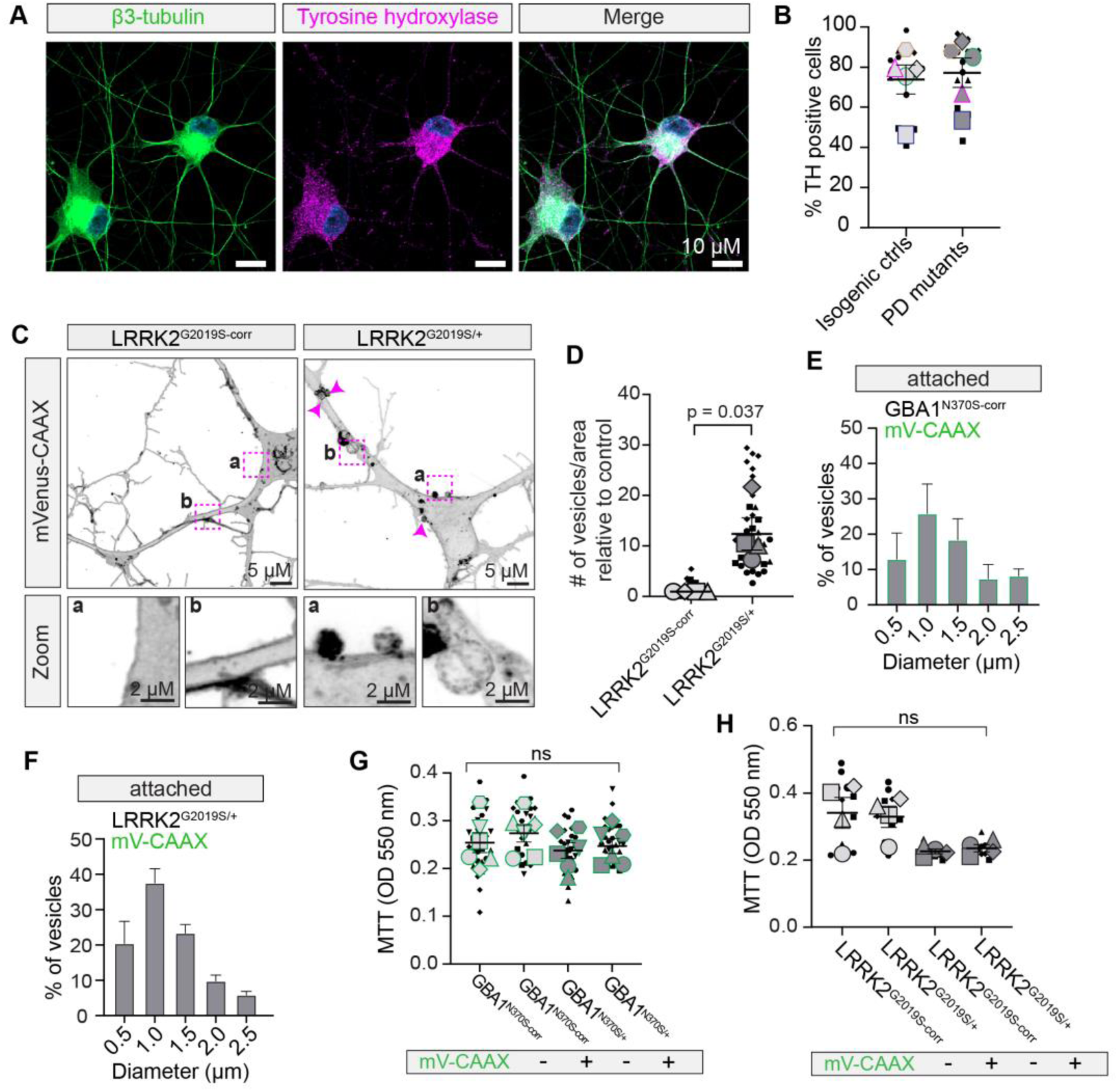
Increased mVenus-CAAX-positive ectosomes from *GBA1* and *LRRK2* patient-derived dopaminergic neurons. **(A)** iPSC-derived dopaminergic neurons fixed and immunostained for β3-tubulin and tyrosine hydroxylase. **(B)** Percentage of tyrosine hydroxylase positive dopaminergic neurons in *GBA1* and *LRRK2* mutant lines and their isogenic controls. Graph is depicted as a superplot where biological replicates are shown in large shapes and technical replicates are shown as small shapes. **(C)** Live-cell maximum intensity projection of *LRRK2-*G2019S and isogenic corrected control (corr) iPSC-derived dopaminergic neurons expressing mVenus-CAAX. Boxed areas highlighting ectosomes forming at the plasma membrane are magnified below. **(D)** Number of mVenus-CAAX positive buds per cell area in *LRRK2-*G2019S and control iPSC-derived dopaminergic neuron. *n* = 3 independent experiments; mean ± SEM; One sample t-test. Graph is depicted as a superplot where biological replicates are shown in large shapes and technical replicates are shown as small shapes. **(E-F)** Size distribution of attached mVenus-CAAX-positive vesicles in *GBA1* and *LRRK2* mutant lines. *n* = 4 independent experiments; mean ± SEM. **(G-H)** Cytotoxicity assessment of mVenus-CAAX expression in *GBA1* and *LRRK2* mutant lines and their isogenic controls with MTT assay. *n* = 4 independent experiments; 3 technical replicates/experiment; mean ± SEM; One-way ANOVA with Dunn’s multiple comparison. Graph is depicted as a superplot where biological replicates are shown in large shapes and technical replicates are shown as small shapes.

**Extended Data Figure 4.**
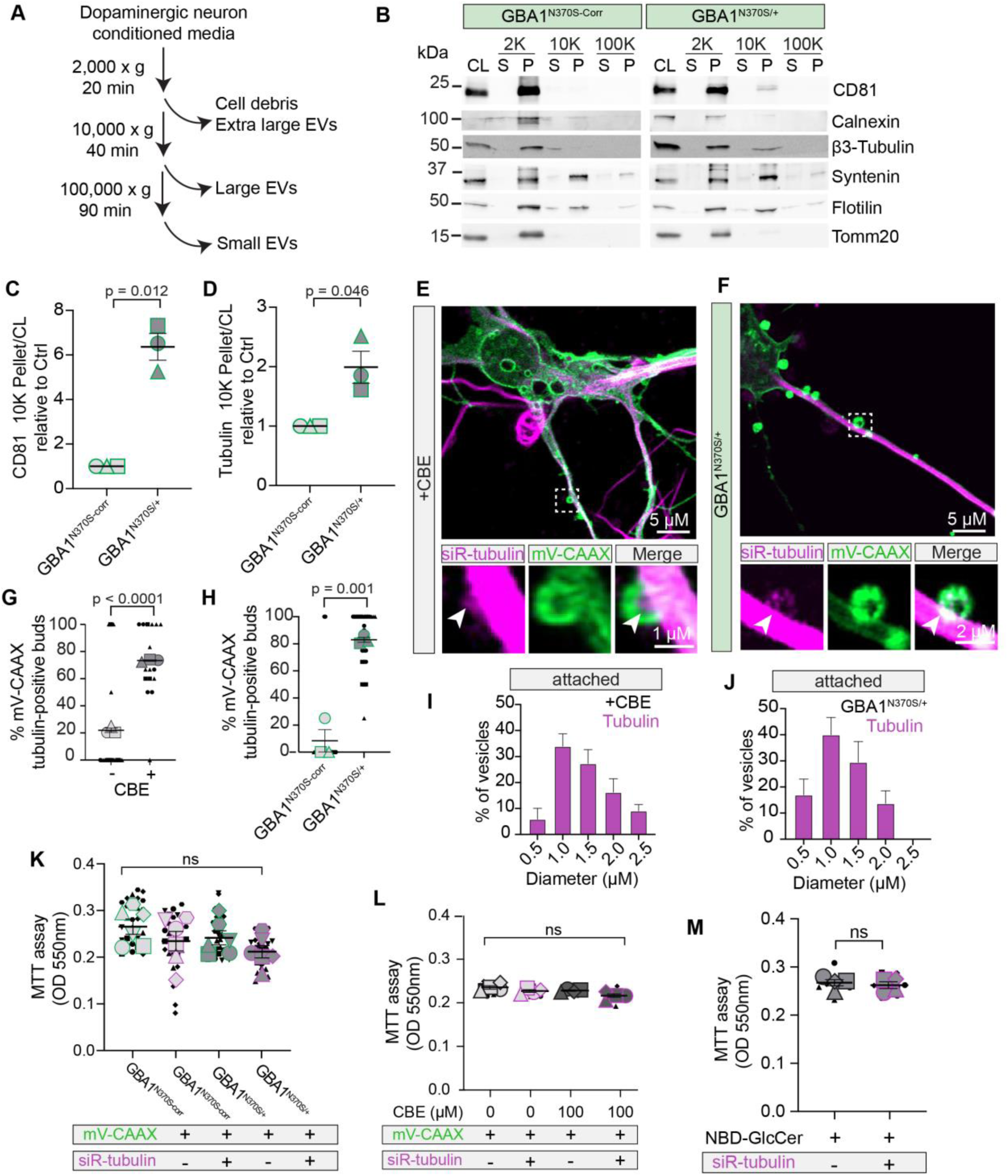
Increased tubulin-positive ectosomes from *GBA1* and *LRRK2* patient-derived dopaminergic neurons. (**A**) Schematic of differential ultracentrifugation procedure used to separate extracellular vesicles (EVs). (**B-D**) Western blotting of centrifugation fractions prepared from the *GBA1*-N370S mutant and its isogenic control as in A. CL, cell lysate; S, supernatant; P, pellet. Quantification CD81 and tubulin present in 10,000 x g fractionated pellet normalized to the cell lysate and relative to the isogenic control. *n* = 3 independent experiments; mean ± SEM; One sample t-test. Graph shows the experimental averages. **(E)** Live-cell image of cortical neurons expressing mVenus-CAAX treated with siR-tubulin and CBE. Boxed areas highlighting an ectosome containing tubulin is magnified below. **(F)** Live-cell image of the *GBA1*-N370S neuron expressing mVenus-CAAX treated with siR-tubulin. Boxed areas highlighting an ectosome containing tubulin is magnified below. (**G-H**) Percentage mVenus-CAAX positive buds containing tubulin induced by CBE and in *GBA1*-N370S neuron. *n* = 3 independent experiments; mean ± SEM; Unpaired t-test. Graph is depicted as a superplot where biological replicates are shown in large shapes and technical replicates are shown as small shapes. (**I-J**) Size distribution of mVenus-CAAX-positive ectosome buds from neurons treated with siR-tubulin and 100 µM CBE or in siR-tubulin treated *GBA1*-N370S neurons. (**K-L**) Cytotoxicity assessment of siR-tubulin treatment on mVenus-CAAX expressing *GBA1*-N370S and control neurons, or CBE treated cortical neurons by MTT assay. *n* = 4 independent experiments; 3 technical replicates/experiment; mean ± SEM; One-way ANOVA with Dunn’s multiple comparison. Graph is depicted as a superplot where biological replicates are shown in large shapes and technical replicates are shown as small shapes. (**M**) Cytotoxicity assessment of NBD-GlcCer +/− siR-tubulin treatment on cortical neurons by MTT assay. *n* = 4 independent experiments; 3 technical replicates/experiment; mean ± SEM; Mann-Whitney test. Graph is depicted as a superplot where biological replicates are shown in large shapes and technical replicates are shown as small shapes.

**Extended Data Figure 5.**
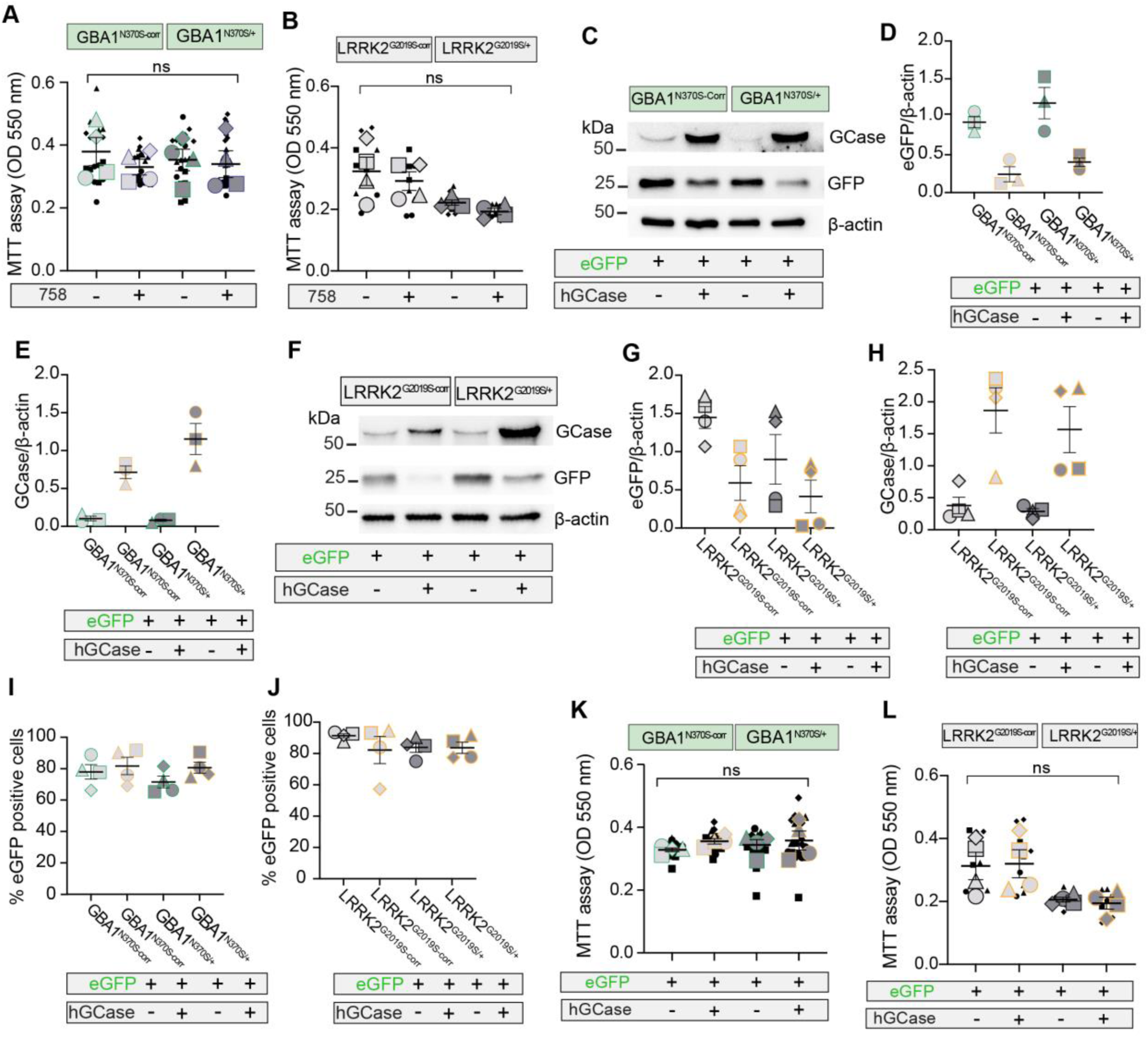
Validation of hGCase/eGFP expression in dopaminergic neurons. (**A-B**) Cytotoxicity assessment of the GCase modulator 758 on *GBA1*-N370S, *LRRK2*-G2019S and corresponding isogenic control neurons by MTT. *n* = 4 independent experiments; 3 technical replicates/experiment; mean ± SEM; One-way ANOVA with Dunn’s multiple comparison. Graph is depicted as a superplot where biological replicates are shown in large shapes and technical replicates are shown as small shapes. (**C-H**) Western blotting of *GBA1*-N370S, *LRRK2*-G2019S and corresponding isogenic control neurons expressing hGCase/eGFP or eGFP and immunoblotted for hGCase, GFP and β-actin. Expression of hGCase and eGFP normalized against β-actin. n=4 independent experiments ; mean ± SEM. Graph shows the experimental averages. (**I-J**) Percentage of eGFP positive dopaminergic neurons in *GBA1*-N370S, *LRRK2*-G2019S and corresponding isogenic controls. *n* = 4 independent experiments; 4 tile scans /2 coverslip/experiment; mean ± SEM. Graph shows the experimental averages. (**K-L**) Cytotoxicity assessment of hGCase/eGFP expression on *GBA1*-N370S, *LRRK2*-G2019S and corresponding isogenic control neurons by MTT. *n* = 4 independent experiments; 3 technical replicates/experiment; mean ± SEM; One-way ANOVA with Dunn’s multiple comparison. Graph is depicted as a superplot where biological replicates are shown in large shapes and technical replicates are shown as small shapes.

**Extended Data Figure 6.**
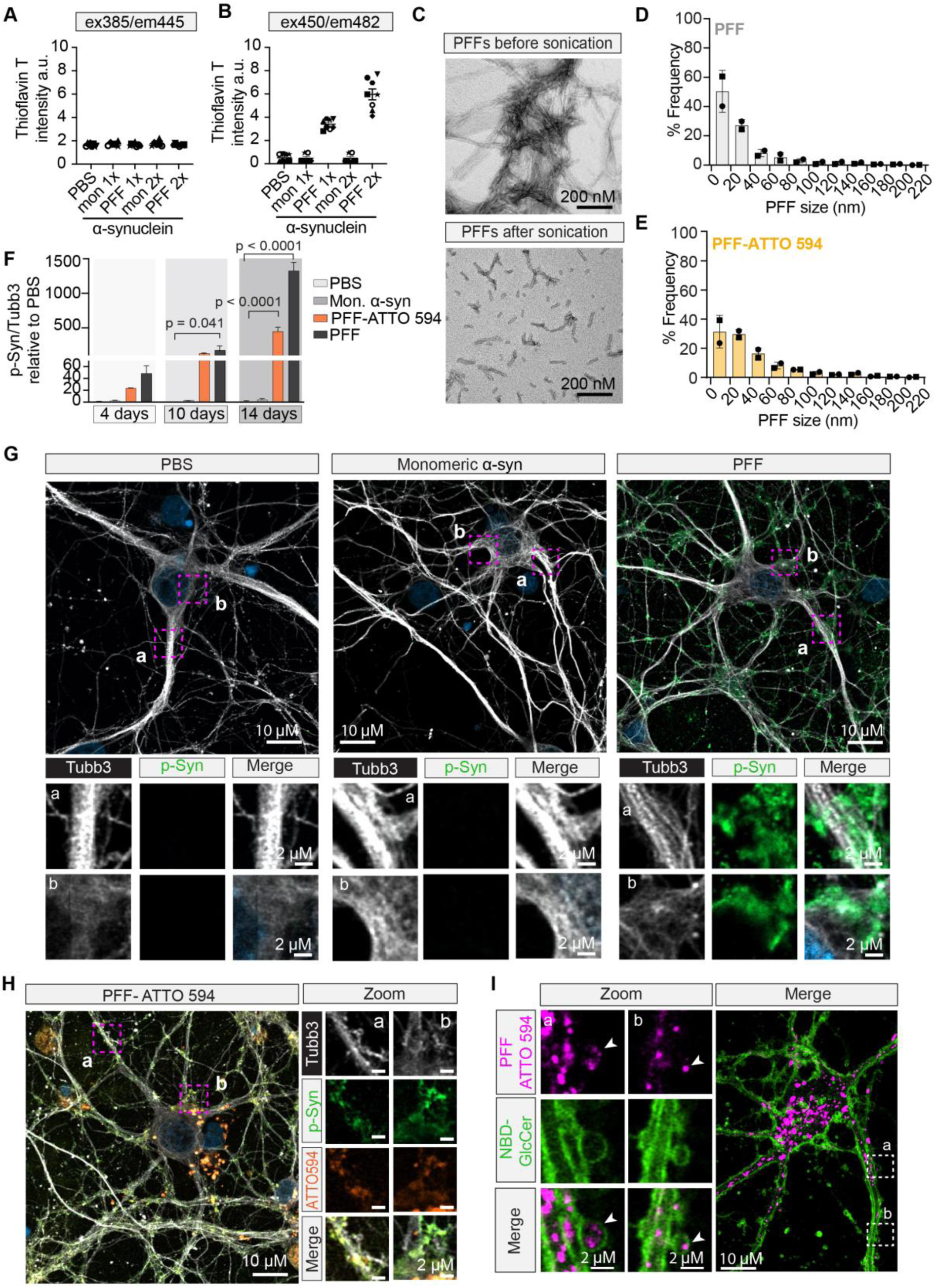
Validation of α-synuclein pre-formed fibrils. (**A-B**) Thioflavin T increases emission at 482 nm with pre-formed fibrils (PFFs) generated from monomeric α-synuclein indicating the presence of beta sheet-rich structures which are absent from monomeric or PBS controls. 2x, double concentration. (**C**) Transmission electron micrographs of α-synuclein PFFs before and after sonication. (**D-E**) Length distribution of fluorescently labelled and unlabeled PFFs after sonication. (**F**) Percent area of pS129-α-synuclein relative to β3-tubulin in primary cortical neurons over time. *n* = 4 independent experiments; mean ± SEM; two-way ANOVA with Dunnett’s multiple comparison test. (**G-H**) Images of cortical cultures at DIV14 treated with PBS, monomeric α-synuclein, tagged PFFs (Atto-594) or untagged PFFs stained for β3-tubulin and pS129-α-synuclein. Boxed areas are magnified below and on the right. (**I**) Live-cell image of ATTO594-α-synuclein fibrils in ectosomes formed upon addition of NBD-GlcCer in cortical neurons. Boxed areas are magnified on the left.

**Extended Data Figure 7.**
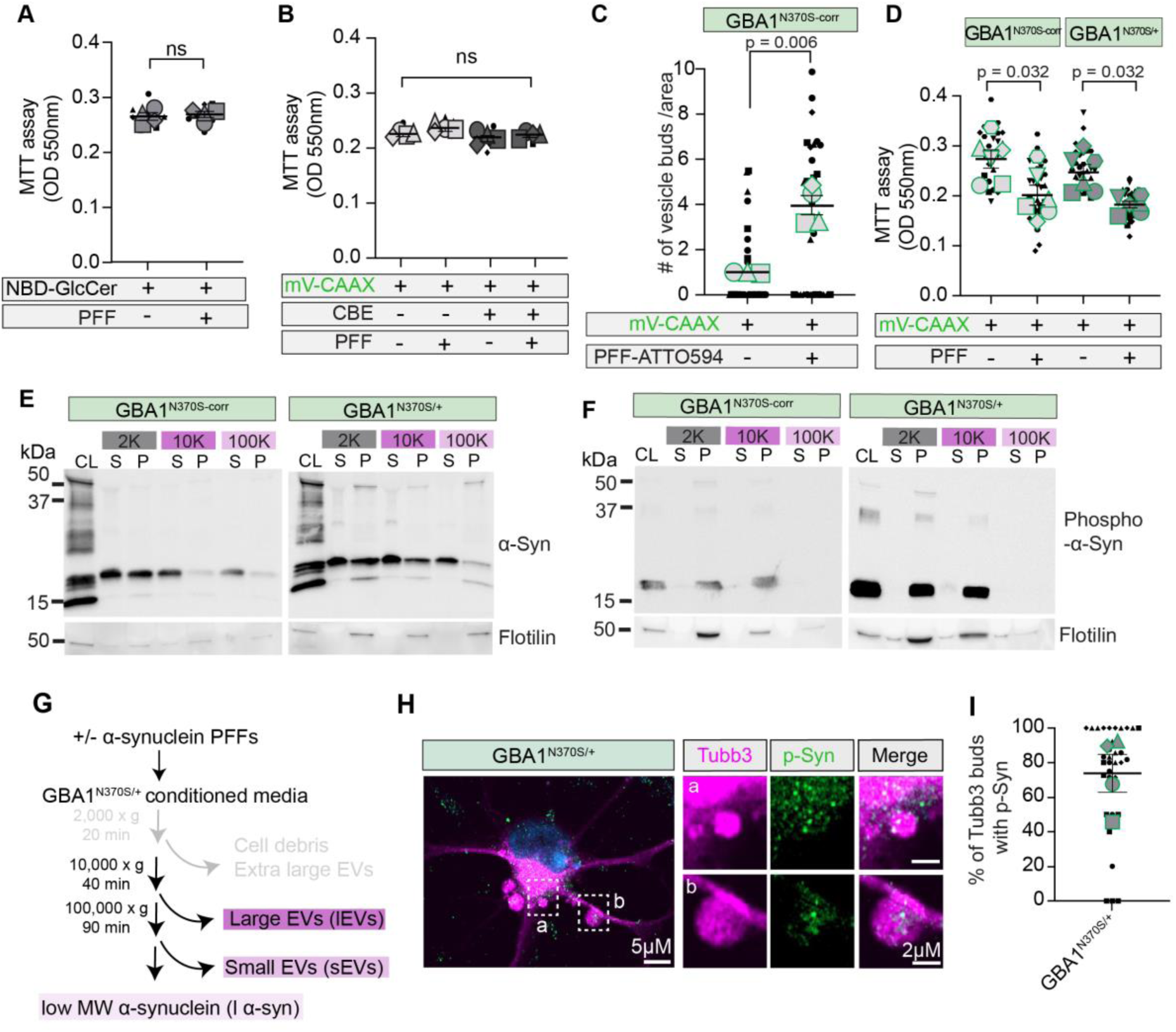
Ectosomes as a vehicle for the transmission of pathogenic α-synuclein. **(A-B)** Cytotoxicity assessment of PFFs on NBD-GlcCer and CBE treated cortical neurons expressing mVenus-CAAX by MTT assay. *n* = 4 independent experiments; 3 technical replicates/experiment; mean ± SEM; Mann-Whitney test and One-way ANOVA with Dunn’s multiple comparison. Graph is depicted as a superplot where biological replicates are shown in large shapes and technical replicates are shown as small shapes. **(C)** Number of mVenus-CAAX positive vesicles per area in *GBA1* isogenic control neurons +/− PFFs. n ≥ 3 independent experiments; 10 cells/coverslip/treatment; mean ± SEM; One sample t-test. Graph is depicted as a superplot where biological replicates are shown in large shapes and technical replicates are shown as small shapes. **(D)** Cytotoxicity assessment of PFFs on mVenus-CAAX expressing *GBA1*-N370S and isogenic control neurons by MTT. *n* = 4 independent experiments; 3 technical replicates/experiment; mean ± SEM; One-way ANOVA with Dunn’s multiple comparison. Graph is depicted as a superplot where biological replicates are shown in large shapes and technical replicates are shown as small shapes. **(E-F)** Western blotting of centrifugation fractions derived from *GBA1*-N370S and control neurons treated with PFFs and immunoblotted for flotilin-1, pS129-α-synuclein and synuclein. CL, cell lysate; S, Supernatant; P, Pellet. 2K, 2000 x g; 10K, 10,000 x g; 100k, 100,000 x g. n=3 independent experiments. **(G)** Procedures used to separate vesicles and low molecular weight α-synuclein released by *GBA1*-N370S neurons by differential centrifugation. **(H**) Image of fixed *GBA1*-N370S dopaminergic showing pS129-α-synuclein in tubulin-positive ectosomal bud. Boxed areas are magnified on the right. **(I)** Percentage of tubulin-positive buds containing pS129-α-synuclein in PFF-treated *GBA1*-N370S dopaminergic. *n* = 4 independent experiments; 10 cells/experiment. Mean ± SEM. Graph is depicted as a superplot where biological replicates are shown in large shapes and technical replicates are shown as small shapes.

**Extended Data Figure 8.**
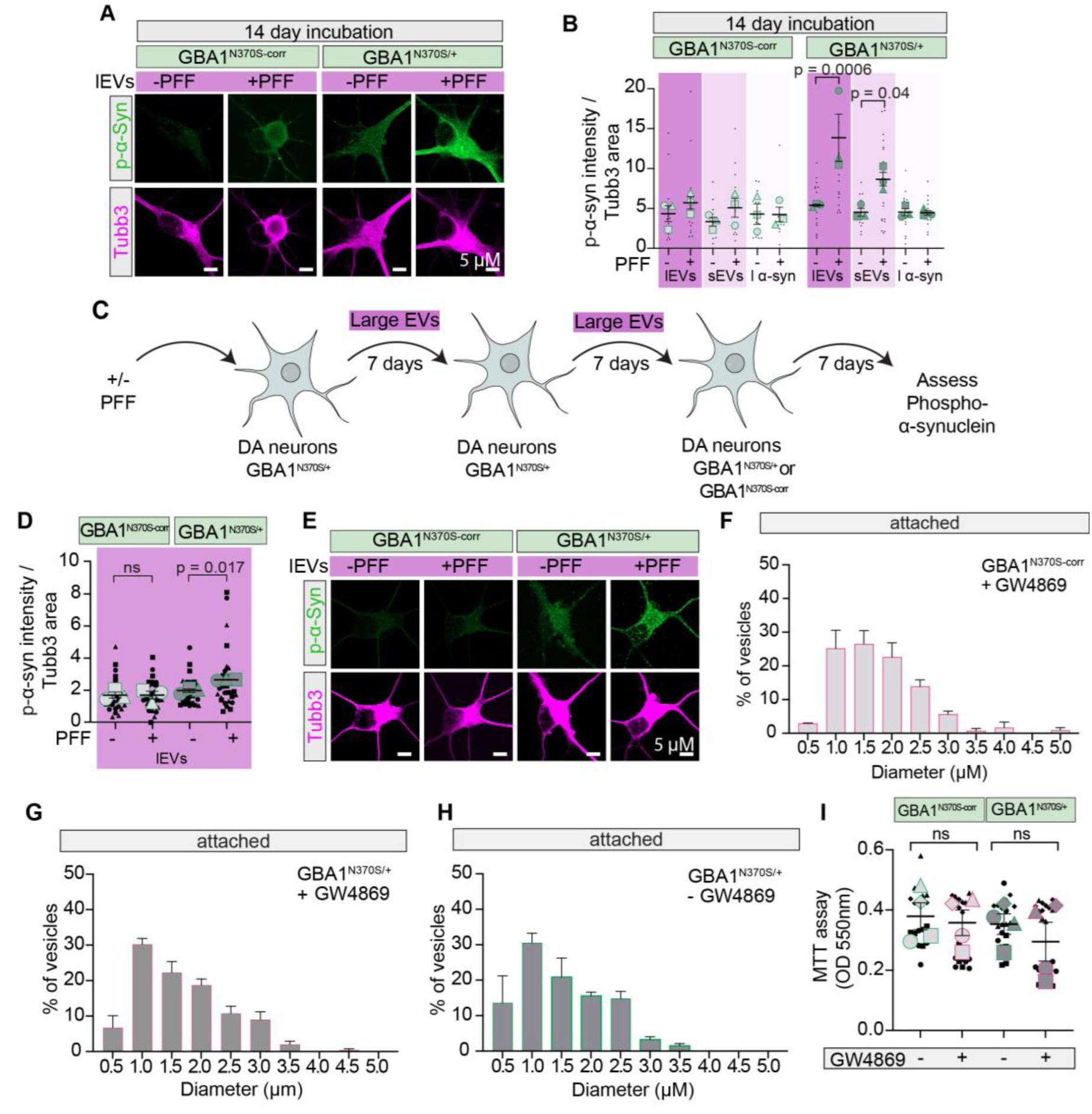
Ectosome-mediated propagation of phosphorylated α-synuclein in *GBA1*-N370S neurons. **(A-B)** Maximum intensity projections of *GBA1*-N370S and control neurons incubated with large vesicle fraction derived from *GBA1*-N370S +/− PFF for 14 days and stained with β3-tubulin and pS129-α-synuclein. Quantifications of the amount of intracellular pS129-α-synuclein normalized to the neuron area in *GBA1*-N370S and control neurons treated with different centrifugation fractions after 14 days of incubation. *n* = 3 independent experiments; 6 cells/experiment. Mean ± SEM; One-way ANOVA with uncorrected Fisher’s LSD. Graph is depicted as a superplot where biological replicates are shown in large shapes and technical replicates are shown as small shapes. **(C)** Schematic representation of the multi-step propagation of pathogenic α-synuclein through large vesicles/ectosomes. (**D-E**) Quantifications of the amount of intracellular pS129-α-synuclein normalized to the neuron area in *GBA1*-N370S and control neurons treated with large vesicle fraction from multi-step process depicted in (C) after 7 days of incubation. *n* = 4 independent experiments; 8 cells/experiment. Mean ± SEM; One-way ANOVA with uncorrected Fisher’s LSD. Graph is depicted as a superplot where biological replicates are shown in large shapes and technical replicates are shown as small shapes. Max intensity projections of *GBA1*-N370S and control neurons incubated with the second-generation large vesicle fractions derived from *GBA1*-N370S. **(F-H)** Size distribution of attached β3-tubulin-positive vesicles from *GBA1-*N370S +/− GW4869 and corresponding isogenic control neurons. *n* = 4 independent experiments; bars show mean ± SEM. **(I)** Cytotoxicity assessment of GW4869 on *GBA1-*N370S and isogenic control neurons by MTT assay. *n* = 4 independent experiments; 3 technical replicates per experiment; bars show mean ± SEM; One-way ANOVA, with Dunn’s multiple correction test. Graph is depicted as a superplot where biological replicates are shown in large shapes and technical replicates are shown as small shapes.

**Supplementary Video 1. Release of tdTomato positive vesicles from neurons in vivo**

A super-resolution microscopy image of a coronal brain section showing detached tdTomato-NBD-GlcCer positive puncta in 3D.

