## Supplementary information iPSC line validation for "Glucosylceramide induced ectosomes propagate pathogenic α-synuclein in Parkinson’s disease"

### GBA L444P-correction/3575

☐ Control line ☒ Disease line
Gene edited? Yes

#### General information

| | |
|---|---|
| Cell line name | GBA L444P-correction/3575 |
| Biosample ID | 3575 |
| Lines from same donor | 3575 |

#### Donor information

| | |
|---|---|
| Sex | MALE |
| Age | 58 YEARS |
| Race | N/A |

#### Culture conditions

| | |
|---|---|
| Coating/ medium | MATRIGEL/mTeSR1 |
| Passage method | Gentle cell dissociation reagent |

#### Derivation

| | |
|---|---|
| Primary cell line | PBMC |
| Reprograming method | EPISOMAL |
| Reprogramming factors | Bcl-Xl ☒ Myc ☒<br>Nanog ☐ SOX2 ☒<br>LIN28 ☐ KLF4 ☒<br>OCT3/4 ☒ |

#### Disease status

| | |
|---|---|
| Disease | Parkinson's Disease |
| Affected gene | GBA |
| Disease mutation/family history | GBA L444P-correction |

#### Genetic modification

| | |
|---|---|
| Modification | CRISPR-Cas9 Knock in |
| Gene | GBA |
| Gene ID | ENSG00000177628 |
| Chromosome location | Chromosome 1: 155,234,452-155,241,249 reverse strand |
| gRNA1 | UGCCAGUCAGAAGAACGACC |
| Delivery method | Lonza Nucleofection |
| Subclone IDs | G5 |
| Description | One gRNA was designed using benchling.com to generate one DSB in exon 10 (ENSE00003644399) of transcript GBA-202 (ENST00000368373.8), 3bp from the target nucleotide, using Cas9 nuclease; the mutation L444P was corrected by HDR using the ssODN template ATTCCTGAGGGCTCCCAGAGAGTGGGGCTGGTTGCCAGTCAGAAGAACGACCTCGACGCAGTGGCACTGATGCATCCCGATGGCTCTGCTGTTGTGGTCGTG<br><br>Edited alleles were detected by ddPCR with primers GCTGGTTGCCAGTCAGAA and GAGAGTGTGATCCTGCCAA, and Affinity probes mutant: /5HEX/ CGAC+C+C+GGAC /3IABkFQ/ and corrected: /56-FAM/ CGA+C+C+TC+G+AC /3IABkFQ/;<br><br>Primers for PCR/Sanger sequencing were GCTGGTTGCCAGTCAGAA and GAGAGTGTGATCCTGCCAA (amplicon: 157bp); |

Sanger DNA sequence of GBA L444P-correction /3575:

GCTGGTTGCCAGTCAGAAGAACGACCTCGACGCAGTGGCACTGATGCATCCCGATGGCTC
TGCTGTTGTGGTCGTGCTAAACCGGTGAGGGCAATGGTGAGGTCTGGGAAGTGGGCTGAA
GACAGCGTTGGGGGCCTTGGCAGGATCACACTCTC

**3575**

**GBA L444P-correction/3575**

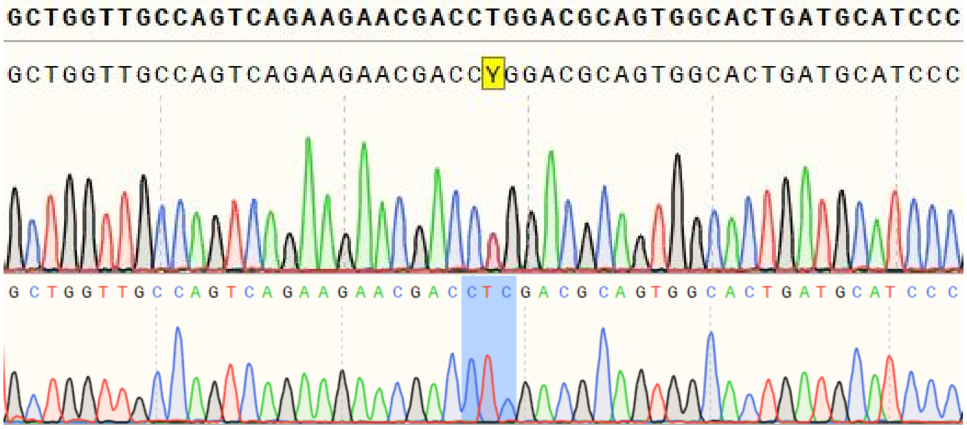


**Figure 1. DNA Sanger sequence of the target codon area in exon 10 of GBA.**
The blue shaded area confirms that the wt codon CTC has replaced the mutant codon CCG, correcting the missense mutation L444P.

#### Characterization:

##### Authentication by STR analysis

| Marker | 3575 | | GBA L444P-correction/3575 | |
|---|---|---|---|---|
| | Allele 1 | Allele 2 | Allele 1 | Allele 2 |
| AMEL | X | Y | X | Y |
| CSF1PO | 10 | 10 | 10 | 10 |
| D13S317 | 11 | 12 | 11 | 12 |
| D16S539 | 9 | 10 | 9 | 10 |
| D21S11 | 29 | 31.2 | 29 | 31.2 |
| D5S818 | 11 | 15 | 11 | 15 |
| D7S820 | 8 | 9 | 8 | 9 |
| TH01 | 9 | 9.3 | 9 | 9.3 |
| TPOX | 8 | 11 | 8 | 11 |
| vWA | 14 | 18 | 14 | 18 |

#### Genotyping analysis:

| | |
|---|---|
| Passage number | P6+C8 |
| Karyotyping | Normal 46, XY |
| qPCR (Genetic Analysis kit) | Normal |

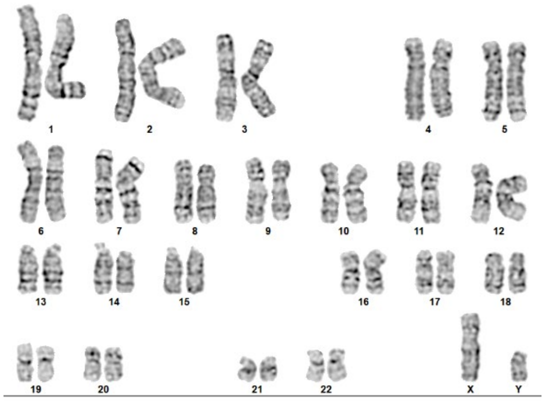


**Figure 2. G-band assay show normal karyotype of parental line GBA L444P-correction/3575, 46, XY.**

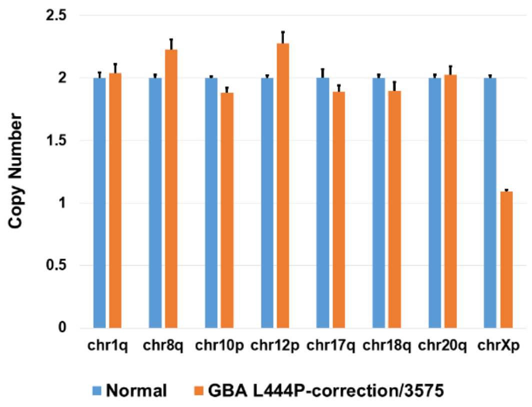


**Figure 3. Genetic stability assay show normal chromosome of GBA L444P-correction/3575.**

#### Pluripotency analysis:

| Marker | Expressed? |
|---|---|
| Nanog | Yes |
| Tra-1-60 | Yes |
| SSEA-4 | Yes |
| OCT3/4 | Yes |

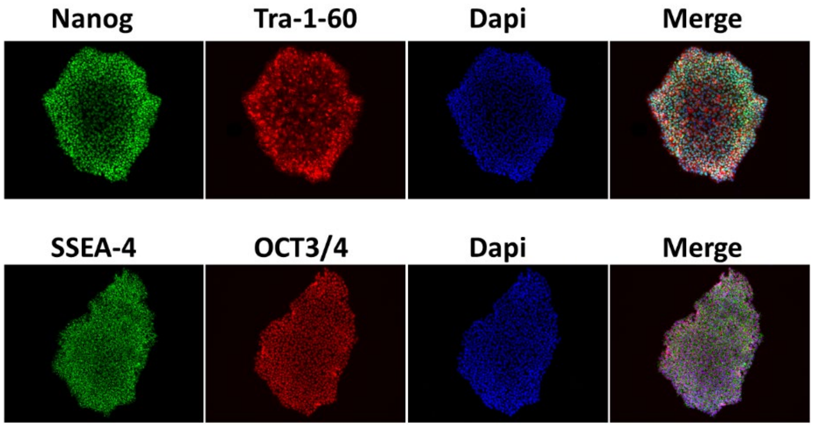


**Figure 4. Immunostaining of pluripotency markers on GBA L444P-correction/3575.**

#### Microbiology/virus screening:

| | |
|---|---|
| Mycoplasma test | Negative |
| Hepatitis B | Negative |
| Hepatitis C | Negative |
| HIV1 or 2 | Negative |

### LRRK2 G2019S-correction/3081

☐ Control line ☒ Disease line
Gene edited? Yes

#### General information

| | |
|---|---|
| Cell line name | LRRK2 G2019S-correction/ 3081 |
| Biosample ID | 3081 |
| Lines from same donor | 3081 |

#### Donor information

| | |
|---|---|
| Sex | FEMALE |
| Age | 62 YEARS |
| Race | Caucasian other than French Canadian |

#### Culture conditions

| | |
|---|---|
| Coating/ medium | Matrigel/mTeSR1 |
| Passage method | Gentle cell dissociation reagent |

#### Derivation

| | |
|---|---|
| Primary cell line | PBMC |
| Reprograming method | EPISOMAL |
| Reprogramming factors | Bcl-Xl ☒ Myc ☒<br>Nanog ☐ SOX2 ☒<br>LIN28 ☐ KLF4 ☒<br>OCT3/4 ☒ |

#### Disease status

| | |
|---|---|
| Disease | Parkinson’s Disease |
| Affected gene | LRRK2 |
| Disease mutation/family history | LRRK2 G2019S |

#### Genetic modification

| | |
|---|---|
| Modification | CRISPR-Cas9 Knock in |
| Gene | LRRK2 |
| Gene ID | ENSG00000188906 |
| Chromosome location | Chromosome 12: 40,196,744-40,369,285 forward strand |
| gRNA1(+strand) | CUCAGUACUGCUGUAGAAUG |
| gRNA2(-strand) | GUCAGCAAUCUUUGCAAUGA |
| Delivery method | Lonza nucleofection |
| Subclone IDs | A6 |
| Description | Two gRNAs were designed using benchling.com to generate two DSBs in exon 41 (ENSE00003681812) of transcript LRRK2-201 (ENST00000298910.12), 20bp upstream and 20bp downstream of the target codon, using Cas9 nuclease; the mutation G2019S was corrected by HDR using the ssODN template GAAACCCCACAATGTGCTGCTTTTCACACTGTATCCCAATGCTGCTATCATTGCAAAGATTGCTGACTACGGCATTGCTCAGTACTGTTGCAGGATGGGCATAAAAACATCAGAGGGCACACCAGGTAGGTGATCAGGTCTGT; corrected alleles were detected by ddPCR with primers GCTGCTTTTCACACTGTATCCC and CTGAGGTCAGTGGTTATCCATC, and Affinity probes mutant: /5HEX/ A+CTA+C+A+G+CA+TTG/3IABkFQ/ and corrected: /56-FAM/A+CTA+C+G+G+CATT/3IABkFQ/; primers for PCR/Sanger sequencing were TGGCAAGTACAACAAAATCCCA and GCCTCACAAGTGCCAACAAT (amplicon: 900bp); |

Sanger DNA sequence (corrected allele):

CTTCTTATTTACATACTTACATTTTGAATAGTTAAAATATTCATATGATCATTGAGAGAATTCAGAATTG
CCTTTAAGTAATTGTTCACATATACAAAAGAAAAGTCTCCAAAAATTGGGTCTTTGCCTGAGATAGATT
TGTCTTAAAATTGAAATCATTCACTTATCAGATTTGACCCTTTTTTAAAGCATAACTTTGCTGTGTAATA
TTAGACTTATATGTTTTGATTTCCTTCTACAATATCTCTTAACTTTAAGGGACAAAGTGAGCACAGAAT
TTTTGATGCTTGACATARTGRACATTTATWTTTAAGGAAATTAGGACAAAAATTATTATAATGTAATCA
CATTTGAATAAGATTTCCTGTGCRTTTTCTGGCAGATACCTCCACTCRRCCRTGATTATRTACCGAGA
CCTGAAACCCCACAATGTGCTGCTTTTCACACTGTATCCCAATGCTGCCATCATTGCAAAGATTGCTG
ACTACGGCRTTGCTCAGTACTGTTGCAGGATGGRCATAAWAACATCAGAKGGCACACCAGGTASGT
GATCAGGTCTGTCTCAYAATTCTATCTTCAGGATGGATAACCACTGACCTCASATGTGAGTTCAGAAG
AGTCAAAGGAARACAKAGTCTATCACATTGTGACAGAGSTTATTTGTGAARMATGCAAGCATCACATG
YGATTTTTATCATGATTTGTAGGAAAAACATGGATGTACTTTCCGCTWACTGAACWTAACGATARAAG
TGWC

Sanger DNA sequence (wt allele):

CTCTATTTACATRCTTACATTTTGAATAGTTAAAATATTCATATGATCATTGAGAGAATTCAGAATTGCC
TTTAAGTRATTGTTCACATATACAAAAGAAAAGTCTCCAAAAAKTGGGTCTTTGCCTGAGATAGATTTG
TCTTAAAATTGAAATCATTCACTTATCAGATTTGACCCTTTTTTAAAGCATAACTTTGCTGTGTAATATT
AGACTTATATGTTTTGATTTCCTTCTACAATATCTCTTAACTTTAAGGGACAAAGTGAGCACAGAATTT
TTGATGCTTGACATARTGRACATTTATWTTTAAGGAAATTAGGACAAAAATTATTATAATGTAATCACA
TTTGAATAAGATTTCCTGTGCRTTTTCTGGCAGATACCTCCACTCRRCCRTGATTATRTACCGAGACC
TGAAACCCCACAATGTGCTGCTTTTCACACTGTATCCCAATGCTGCCATCATTGCAAAGATTGCTGAC
TACGGCRTTGCTCAGTACTGCTGTAGAATGGRGATMAAAACATCAGAKGGCACACCAGGTASGTGAT
CAGGTCTGTCTCAYAATTCTATCTTCAGGATGGATAACCACTGACCTCASATGTGAGTTCAGAAGAGT
CARAAGGAARACAGAGTCTATCACATTGTGACAGAGSTYTATTTGTGARMATGCAGCAKCACATGYG
ATTTTAKCATGAATYTGGTWGAAMAACATGATGTACTTTTCMGCTAWACKGAACTAWACGATGA

A

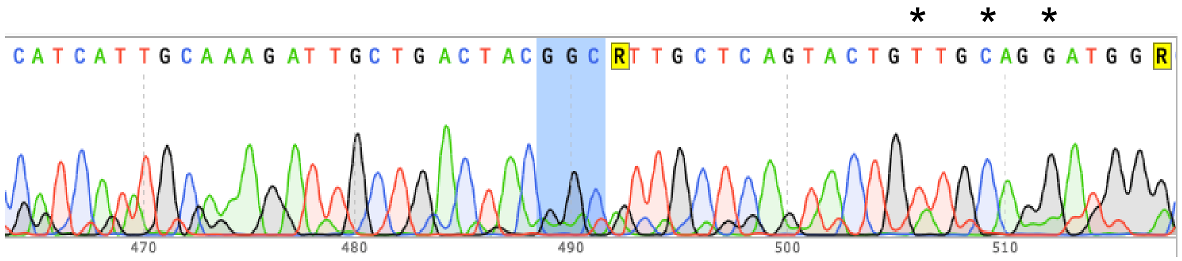


B

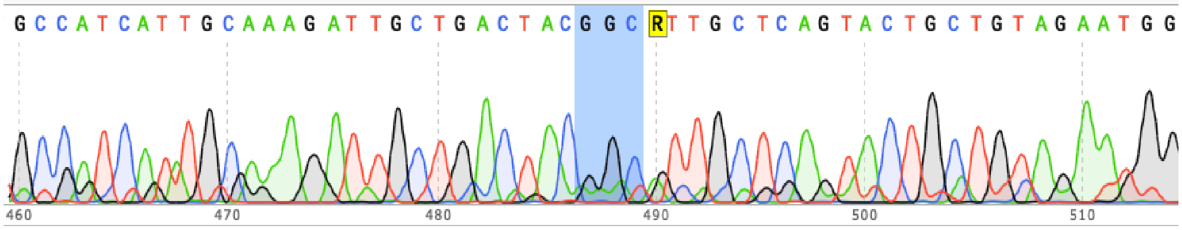


**Figure 1. DNA Sanger sequence of the target codon area in exon 41 of LRRK2.**

(A) The blue shaded area confirms that the wt codon GGC has replaced the mutant codon AGC, correcting the missense mutation G2019S. *Note that three silent mutations C>T, T>C and A>G were introduced in the corrected allele (heterozygous), 15, 18, and 21 bp downstream of the target codon, respectively. (B) The wt allele remained unchanged.

#### Characterization:

##### Authentication by STR analysis

| | 3081 | | LRRK2 G2019S-correction/3081 | |
|---|---|---|---|---|
| Marker | Allele 1 | Allele 2 | Allele 1 | Allele 2 |
| AMEL | X | X | X | X |
| CSF1PO | 7 | 12 | 7 | 12 |
| D13S317 | 9 | 11 | 9 | 11 |
| D16S539 | 11 | 11 | 11 | 11 |
| D21S11 | 28 | 31.2 | 28 | 31.2 |
| D5S818 | 12 | 12 | 12 | 12 |
| D7S820 | 8 | 10 | 8 | 10 |
| TH01 | 7 | 9.3 | 7 | 9.3 |
| TPOX | 8 | 8 | 8 | 8 |
| vWA | 15 | 18 | 15 | 18 |

##### Genotyping analysis:

| | |
|---|---|
| Passage number | P8+C4 |
| Karyotyping | Normal 46, XX |
| qPCR (Genetic Analysis kit) | Duplication of chr20q |

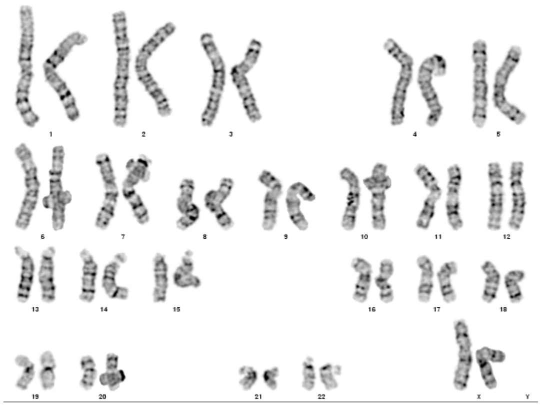


**Figure 2.G-band assay show normal karyotype of LRRK2 G2019S-correction/3081, 46, XX.**

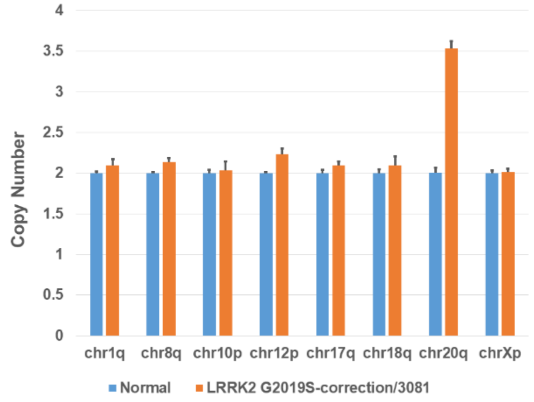


**Figure 3. Genetic stability assay show duplication of chr20q on LRRK2 G2019S-correction/3081.**

**Pluripotency analysis:**

| Marker | Expressed? |
|---|---|
| Nanog | Yes |
| Tra-1-60 | Yes |
| SSEA-4 | Yes |
| OCT3/4 | Yes |

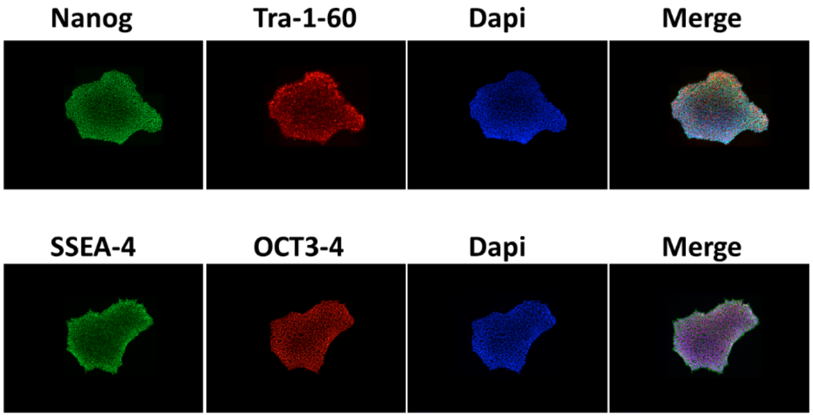


**Figure 4. Immunostaining of pluripotency markers on LRRK2 G2019S-correction/3081.**

**Microbiology/virus screening:**

| | |
|---|---|
| Mycoplasma test | Negative |
| Hepatitis B | Negative |
| Hepatitis C | Negative |
| HIV1 or 2 | Negative |

#### LRRK2 R1441H-correction/3439

☐ Control line ☒ Disease line
Gene edited? Yes

##### General information

| | |
|---|---|
| Cell line name | LRRK2 R1441H-correction/3439 |
| Biosample ID | 3439 |
| Lines from same donor | 3439 |

##### Donor information

| | |
|---|---|
| Sex | MALE |
| Age | 54 YEARS |
| Race | Creole Haitian |

##### Culture conditions

| | |
|---|---|
| Coating/ medium | MATRIGEL/mTeSR1 |
| Passage method | Gentle cell dissociation reagent |

##### Derivation

| | |
|---|---|
| Primary cell line | PBMC |
| Reprograming method | EPISOMAL |
| Reprogramming factors | Bcl-Xl ☒ Myc ☒<br>Nanog ☐ SOX2 ☒<br>LIN28 ☐ KLF4 ☒<br>OCT3/4 ☒ |

##### Disease status

| | |
|---|---|
| Disease | Parkinson's Disease |
| Affected gene | LRRK2 |
| Disease mutation/family history | R1441H |

##### Genetic modification

| | |
|---|---|
| Modification | CRISPR-Cas9 Knock in |
| Gene | LRRK2 |
| Gene ID | ENSG00000188906 |
| Chromosome location | Chromosome 12: 40,196,744-40,369,285 forward strand. |
| gRNA1 | AAGAAGAAGCGTGAGCCTGG |
| Delivery method | Lonza Nucleofection |
| Description | gRNA was designed using benchling.com to generate one DSB in LRRK2 gene (ENSG00000188906) with Cas9 nuclease; The Corrected cell line was created by HDR using the ssODN template AATGTGAGCAGGCCCAGTTTGAAAGCAAACACAAGAGGGTTTTGTGTCTTTCTCTCCAGGCCCGGGCCTCCTCTTCCCCTGTGATTCTCGTTGGCACACATTTGGATGTT. Edited alleles were detected by ddPCR with primers: LRRK2 1441 tag F: AGTGAATGTCACGGAAAGCA. LRRK2 1441 tag R: CGCTTATTCAGGAGTTCCTTGG. LRRK2 1441R Mut probe: 56-FAM/ CC+G G+GC +CT+C CT /3IABkFQ/. SYT1-Ftaq: AGC CAT AGT CGC AGT CCT. SYT1-Rtaq: ACC TGA TCT TTC ATC GTC TTC C. SYT1-wtHEX: /5HEX/ AA+GAA+GAA+G+G+GA/3IABkFQ/. Primers for PCR/Sanger sequencing were forward primer: AGAATTGGGTTAAGAAAGGCCA and reverse primer: TCCTCGGTGGCATTCACAAA (amplicon:385 bp); sequenced with primer AGAATTGGGTTAAGAAAGGCCA and TCCTCGGTGGCATTCACAAA |

Sanger DNA sequence of LRRK2 R1441H-correction/3439

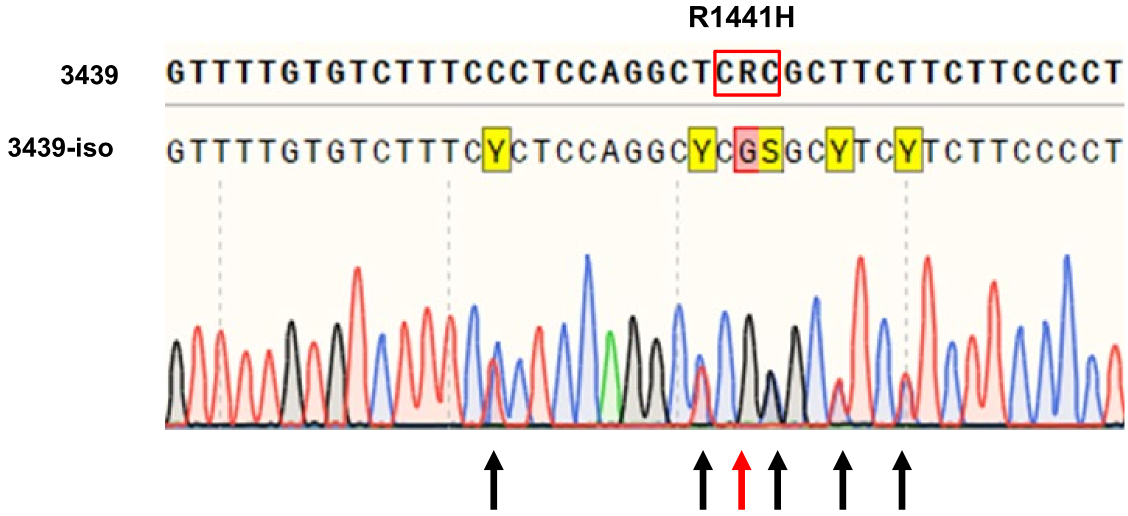


**Figure 1. DNA Sanger sequence of the target codon area of LRRK2 R1441H-correction/3439.**

The red square area confirms that the mutation codon CRC (His) has replaced the mutant codon CGC (Arg) (red arrow), correcting the missense mutation R1441H. Note that five silent mutations were introduced in the corrected allele (black arrows).

#### Characterization:

##### Authentication:

| Marker | LRRK2 R1441H-correction/3439 | | 3439 | |
|---|---|---|---|---|
| | Allele 1 | Allele 2 | Allele 1 | Allele 2 |
| AMEL | X | Y | X | Y |
| CSF1PO | 12 | 13 | 12 | 13 |
| D13S317 | 8 | 8 | 8 | 8 |
| D16S539 | 10 | 13 | 10 | 13 |
| D21S11 | 29 | 30 | 29 | 30 |
| D5S818 | 10 | 11 | 10 | 11 |
| D7S820 | 11 | 12 | 11 | 12 |
| TH01 | 8 | 10 | 8 | 10 |
| TPOX | 8 | 8 | 8 | 8 |
| vWA | 17 | 17 | 17 | 17 |

#### Genotyping analysis:

| | |
|---|---|
| Passage number | P6+C11 |
| Karyotyping | Normal 46, XY |
| qPCR (Genetic Analysis kit) | Normal |

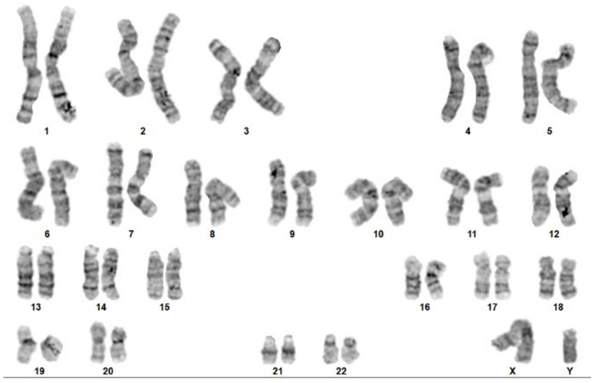

**Figure 2. G-band assay show normal karyotype of LRRK2 R1441H-correction/3439, 46, XY.**

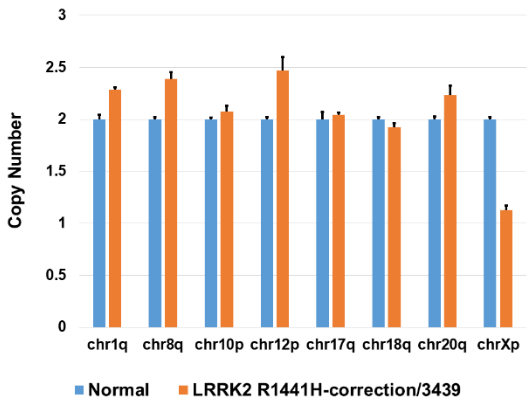


**Figure 3. Genetic stability assay show normal chromosome of LRRK2 R1441H-correction/ 3439.**

#### Pluripotency analysis:

| Marker | Expressed? |
|---|---|
| Nanog | Yes |
| Tra-1-60 | Yes |
| SSEA-4 | Yes |
| OCT3/4 | Yes |

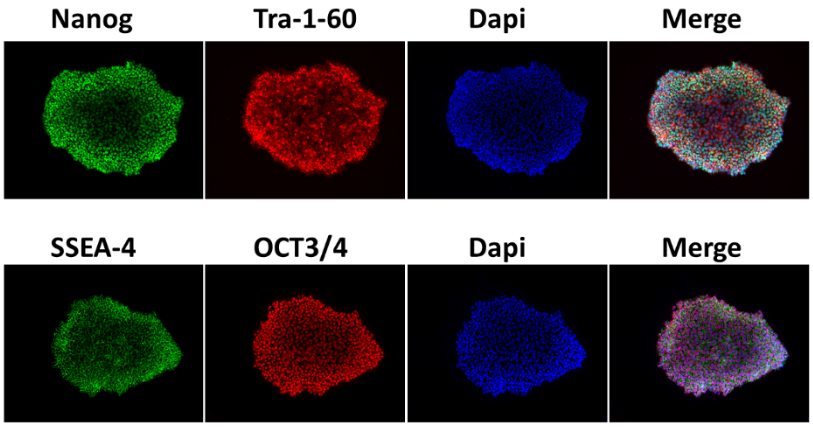


**Figure 4. Immunostaining of pluripotency markers on LRRK2 R1441H-correction/3439.**

#### Microbiology/virus screening:

| | |
|---|---|
| Mycoplasma test | Negative |
| Hepatitis B | Negative |
| Hepatitis C | Negative |
| HIV1 or 2 | Negative |
